# Comparison of seven modelling algorithms for GABA-edited ^1^H-MRS

**DOI:** 10.1101/2021.11.15.468534

**Authors:** Alexander R. Craven, Pallab K. Bhattacharyya, William T. Clarke, Ulrike Dydak, Richard A. E. Edden, Lars Ersland, Pravat K. Mandal, Mark Mikkelsen, James B. Murdoch, Jamie Near, Reuben Rideaux, Deepika Shukla, Min Wang, Martin Wilson, Helge Zöllner, Kenneth Hugdahl, Georg Oeltzschner

## Abstract

Edited MRS sequences are widely used for studying GABA in the human brain. Several algorithms are available for modelling these data, deriving metabolite concentration estimates through peak fitting or a linear combination of basis spectra. The present study compares seven such algorithms, using data obtained in a large multi-site study.

GABA-edited (GABA+, TE = 68 ms MEGA-PRESS) data from 222 subjects at 20 sites were processed via a standardised pipeline, before modelling with FSL-MRS, Gannet, AMARES, QUEST, LCModel, Osprey and Tarquin, using standardised vendor-specific basis sets (for GE, Philips and Siemens) where appropriate.

After referencing metabolite estimates (to water or creatine), systematic differences in scale were observed between datasets acquired on different vendors’ hardware, presenting across algorithms. Scale differences across algorithms were also observed.

Using the correlation between metabolite estimates and voxel tissue fraction as a benchmark, most algorithms were found to be similarly effective in detecting differences in GABA+. An inter-class correlation across all algorithms showed single-rater consistency for GABA+ estimates of around 0.38, indicating moderate agreement. Upon inclusion of a basis set component explicitly modelling the macromolecule signal underlying the observed 3.0 ppm GABA peaks, single-rater consistency improved to 0.44. Correlation between discrete pairs of algorithms varied, and was concerningly weak in some cases.

Our findings highlight the need for consensus on appropriate modelling parameters across different algorithms, and for detailed reporting of the parameters adopted in individual studies to ensure reproducibility and meaningful comparison of outcomes between different studies.

**Highlights:**

- GABA-edited MRS data from 222 healthy adults across 20 research sites were analyzed
- Data were modelled using seven different algorithms, yielding GABA+ and Glx estimates
- Moderate agreement was seen across all the tested algorithms
- Adding a component to represent co-edited macromolecule signals improved concordance
- Baseline modelling emerged as major factor differentiating outcomes

## 1 Introduction

Several software packages and modelling algorithms are available for processing and quantifying MR spectroscopy (MRS) data. While they are all designed to extract quantitative estimates of metabolite levels from spectra, the packages differ significantly in their approach to processing and modelling the underlying data, and isolating the components of interest from any artefactual signals therein. This may give rise to systematic differences in metabolite estimates between different software packages. While an effect of “choice of software” has been documented for short-echo-time data ^1–4^, similar studies for GABA-edited MRS quantification are lacking.

Spectral editing experiments ^5,6^, such as the widely used MEGA-PRESS for the selective detection of GABA, present a special case for quantification. In a typical MEGA-PRESS editing sequence, two interleaved sub-spectra are acquired: the edit-ON sub-spectrum in which coupling to GABA spins at 3 ppm is refocused, and the edit-OFF sub-spectrum in which it is not. Subtracting the edit-ON and edit-OFF sub-spectra yields a relatively sparse difference spectrum, featuring prominent broad signal for GABA (with underlying macromolecule contributions) at 3 ppm and co-edited signals including glutamate (Glu) and glutamine (Gln) peaks (usually reported collectively as Glx) around 3.75 ppm, and strong negative peaks close to the editing frequency (primarily N-acetylaspartate (NAA) and N-acetylaspartylglutamate (NAAG)).

Most notable among the challenges for modelling edited spectra are co-edited macromolecular signals coupled to spins near the editing frequency ^6^, some of which appear in the same frequency range as the GABA and Glx signals and therefore interfere with their unambiguous modelling. As they are broad and poorly characterized, no consensus currently exists on how they should be accounted for in the modelling stage. Constrained by the inability to reliably separate GABA and macromolecules, their composite (GABA+) is commonly reported.

A rigorous assessment of the comparability of GABA+ estimates obtained across a range of different analysis software packages is currently lacking. Several prior studies, including ^7–10^, have investigated test-retest reproducibility of GABA+ estimates using a small selection of available software packages, but without detailed examination of the differences in estimates arising between software packages. Each considered data from a single site only. Another study ^11^ has investigated GABA+ estimates from Gannet and Tarquin compared to a simulated “ground truth”, specifically with respect to the influence of signal-to-noise ratio (SNR) and linewidth on estimates, showing that the two algorithms agreed under favourable conditions of linewidth and signal-to-noise ratio but diverged under poorer conditions – however, only two algorithms were included in this analysis. A recent conference paper ^12^ has reported early findings from GABA-edited MEGA-PRESS data showing moderate associations between five different algorithms, with data from four sites (representing two scanner vendors), albeit with divergent processing. A more thorough examination, covering a broader range of sites and an extended selection of contemporary algorithms is required to better characterise the noted discrepancies.

Therefore, to establish the degree to which different software packages agree in estimating GABA+ from MEGA-PRESS data, this study compares GABA+ estimates from seven modelling algorithms: FSL-MRS ^13^, Gannet ^14^, LCModel ^15^, Osprey ^16^, Tarquin ^17,18^, AMARES ^19^, and QUEST ^20,21^, with the last two implemented in the jMRUI software package ^22,23^. Estimates of Glx from the difference spectra are also considered. Detailed characterisation of differences observable across algorithms is essential for meaningful comparison of findings reported from different tools, and particularly in reconciling any discrepancies therein.

## 2 Methods

### 2.1 Data

Data from twenty 3T MRI scanners from the three major manufacturers (GE, Philips, Siemens), each at a different site, were obtained from the Big GABA ^24,25^ repository on NITRC, https://www.nitrc.org/projects/biggaba. GABA-edited spectra (TR/TE = 2000/68 ms, 320 averages, editing at 1.9/7.46 ppm for edit-ON/-OFF, respectively) and corresponding water-unsuppressed reference data (8 or 16 averages) were obtained from a 3 x 3 x 3 cm^3^ voxel in the posterior cingulate region, from 222 consenting adult volunteers (18-36 years of age, approximately even female/male split, having no known neurological or psychiatric illness), in accordance with ethical standards of their respective local institutional review boards (IRB). Subjects consented to the sharing of anonymized data, with allowance for further study. Datasets also included Ti-weighted structural MR images, which were used for tissue segmentation.

This extensive collection of datasets was acquired in an international collaborative study; several aspects have been previously reported ^24–26^, with a focus on comparability across sites and vendors. Full details on the acquisition protocol, software and hardware configurations and sample composition may be found in these papers, and are summarised in Supplementary Table 1. Detailed vendor-specific parameters have been reported previously ^25,27^.

### 2.2 Processing

To the maximum extent practical, data were prepared for each algorithm using a common pipeline, to avoid variations in processing that might otherwise confound observations regarding the model fit. Original data in vendor-specific format were imported using the GannetLoad function from Gannet (v3.1). The GannetLoad function was chosen due to it having the broadest support for the diverse file formats and sequence implementations present in these datasets. This function applies coil combination where necessary, and initial categorisation of individual FIDs into edit-ON/OFF sub-spectra and water reference spectra. Although GannetLoad also incorporates a full processing pipeline, we did not make use of this, instead electing to implement a generalised pipeline in accordance with current consensus recommendations ^28^, using the processing tools from FID-A ^29^. The rationale for this was to provide a common, neutral starting point for quantification across all the algorithms to be assessed, rather than one which may have been tuned for a particular quantification algorithm. Additionally, the standard Gannet pipeline performs line-broadening and zero-filling, which invalidates assumptions for error calculations in linear-combination modelling algorithms such as LCModel.

Initially, motion-corrupted transients were removed by comparing the root mean square of the difference between each transient and the median of transients in the time domain, rejecting those which differed from the mean by more than four standard deviations. This unlikeness metric was calculated independently within the edit-ON and edit-OFF sub-spectra but applied pairwise. To correct for frequency and phase drift, the remaining FIDs were then aligned in the frequency domain using the spectral registration method ^30^, iteratively on a variable, restricted frequency range (~1.6-5.5 ppm in the first iteration, reducing to ~1.8-4.0 ppm in subsequent iterations), before averaging within the edit-ON and edit-OFF sub-spectra. Eddy-current correction (ECC) was applied ^31^, before zero-order phase adjustment of each sub-spectrum according to a dual-Lorentzian model for creatine and choline defined in Gannet ^14^ and implemented in Osprey. Thereafter, edit-ON and edit-OFF sub-spectra were aligned by spectral registration ^30^, and sum and difference spectra were calculated arithmetically on the time domain data, dividing by the number of sub-spectra. All resultant spectra and sub-spectra were frequency-shifted such that the main creatine peak (from the dual-Lorentzian model for creatine and choline) in the sum spectrum appeared at 3.027 ppm. After calculation of the difference spectrum, residual water was filtered from edit-ON and edit-OFF data separately with the HSVD method ^32^. A final automated check was performed to ensure correct ON/OFF ordering and orientation of all resultant spectra, flipping where necessary. Processed data were exported with the same resolution (number of samples and sweep width/dwell time) as the incoming data, without line-broadening or zero-filling. Processing and modelling workflow are summarised in Figure 1.

**Figure 1:**
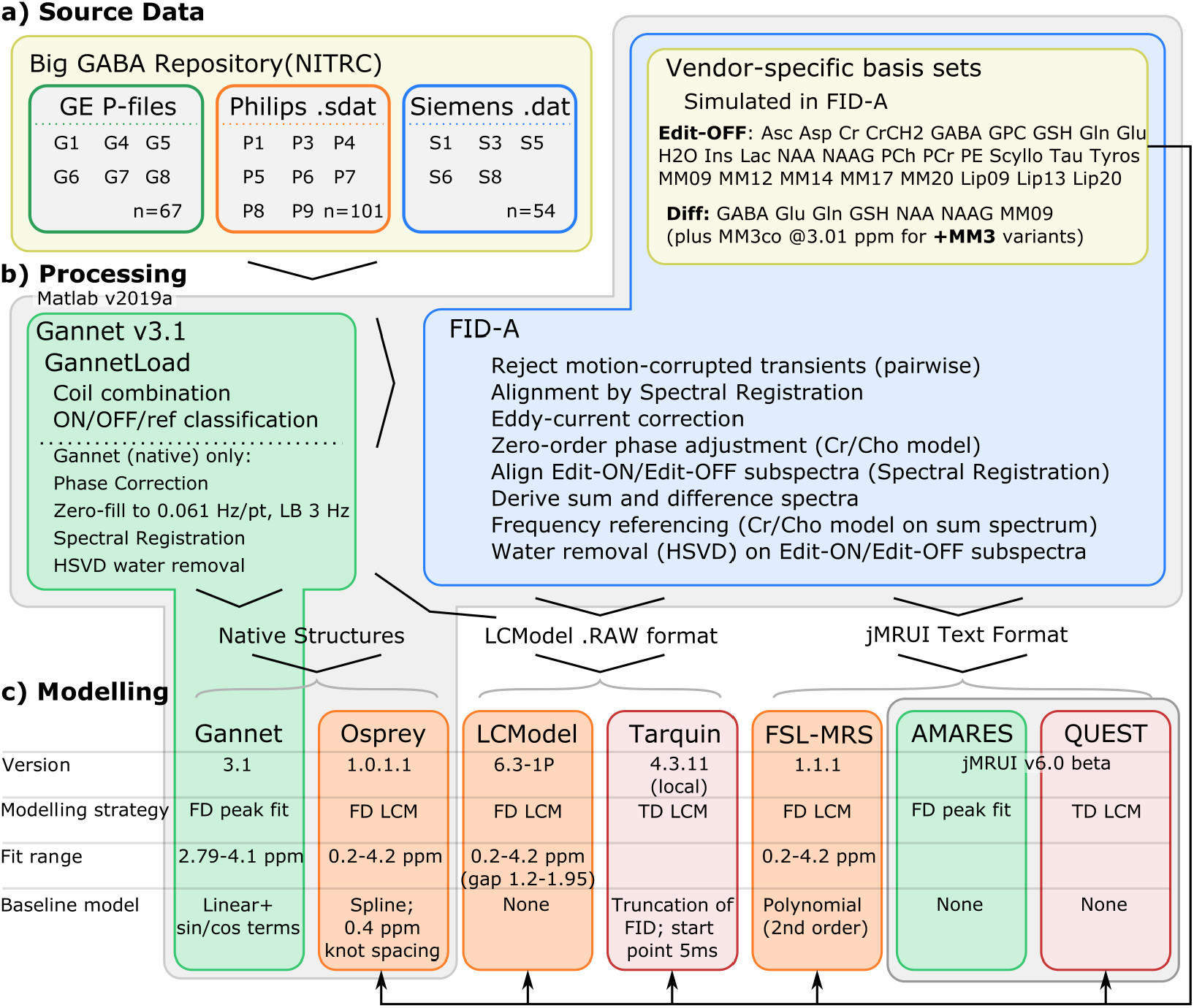
Processing (b) and modelling (c) workflow, summarising key differences between the algorithms assessed.

#### 2.2.1 Quality Control: Processing

Processed spectra were tested against two rejection criteria, designated **R1** and **R2** in subsequent usage:

- R1 captures spectra having strongly aberrant features in the fit range: processing was deemed to have failed if the 0-lag cross-correlation of the normalized, reconstructed frequency domain difference spectrum in the metabolite range (2.6-4.2 ppm) with the normalized mean of all other difference spectra was below 0.5 or differed from the group mean by more than three standard deviations.
- R2 establishes thresholds on basic signal quality metrics: SNR (< 80, defined by maximum peak height around NAAdiff in the [1.8, 2.2] ppm interval, over standard deviation over the [-2, 0] ppm range) and linewidth (FWHM > 10 Hz, ^33^) measured from NAAdiff.

Data deemed to have failed at the processing stage were still passed to the fit algorithms but flagged as having failed and excluded from evaluation of groupwise statistics (such as median estimates) in further analysis.

### 2.3 Initial Fit and Quantification

Identically processed data were fed into each algorithm. To the maximum extent practical, data were modelled using the developer-supplied default or recommended configuration parameters for GABA-edited MEGA-PRESS data, to yield outcomes representative of those which researchers could expect without extensive local optimisation.

Batch processing for all algorithms was automated in Matlab (v2019a), with the exception of the jMRUI-based algorithms for which processed data were exported then processed as batches (grouped by manufacturer and spectral resolution) in a standardised but manual procedure through the jMRUI user interface. As the commonly used default processing pipeline for Gannet incorporates zero-fill and line-broadening factors not present in the standardised pipeline adopted here, we report outcomes both from the standardised processing pipeline (hereafter denoted “Gannet”), and from data processed with Gannet’s own default pipeline, denoted “Gannet (native)”. Tarquin fitting is often performed with an internally simulated basis set; we also assess outcomes from this mode of operation, hereafter denoted “Tarquin (internal)”.

Full details on the operation of each method are supplied in the Supplementary Material, section C for quantification of the edited difference spectra. To facilitate concentration scaling to an internal creatine reference, corresponding edit-OFF sub-spectra are also modelled; this is described in the Supplementary Material, section E. Concentration estimates are reported both relative to total creatine (tCr), and with respect to an internal water reference; the complexities and relative merits of each approach are described in ^28^.

As specifics of each algorithm’s water referencing procedure varied considerably, scaling as documented for the respective algorithms was first reversed to yield a raw ratio of signal intensities, before applying tissue-class correction ^34^ using previously derived tissue fractions ^24^. Full details on the adjustment for each algorithm are provided in the Supplementary Material, section D. Water-scaled, tissue-class corrected molar concentration estimates are hereafter denoted “/H_2_O”; concentration estimates scaled to water with no adjustment for tissue class (assuming pure water concentration per eq(3) of ^35^) are also calculated, denoted “/H_2_O_noTC_”.

#### 2.3.1 Basis Set Preparation and Prior Knowledge

All the algorithms examined require some degree of prior knowledge to describe expected spectral features, either in the form of parameter constraints or simulated basis sets. In the present study, prior knowledge was standardised as far as possible: all algorithms requiring a basis set were supplied with the same simulated basis set appropriate to the dataset, whilst both algorithms parameterising individual peaks (Gannet and AMARES) were supplied with similar model parameters.

For comparison of the basis set algorithms (FSL-MRS, LCModel, Osprey, QUEST and Tarquin), a standard simulated basis set specific to each hardware vendor was adopted. As a starting point, vendor-specific basis sets for GABA-edited MEGA-PRESS (TE = 68 ms) that are distributed with Osprey were used; these are derived from fast spatially resolved 2D density-matrix simulations ^36^ implemented in FID-A using ideal excitation pulses and vendor-specific refocusing pulses and timings, and using chemical shifts and J-coupling coefficients from Kaiser et al. ^37^. These incorporated metabolite basis functions for GABA, Glu, Gln, glutathione (GSH), NAA, NAAG and a Gaussian component (FWHM = 10.9 Hz) representing co-edited macromolecules around 0.91 ppm (MM09ex).

A variation of this basis set was created, incorporating an additional Gaussian component at 3.0 ppm (simulated with FWHM = 14 Hz and scaled intensity equivalent to two protons) to represent co-edited macromolecule signal underlying the GABA peak around 3.0 ppm. This component, denoted MM3co, allowed the influence of macromolecule modelling on the various algorithms to be examined; subsequent use of this basis set is annotated with “+MM3”. The interaction of this component with baseline stiffness and soft constraint models similar to those of ^38–40^ is explored for Osprey and LCModel in a supplementary analysis, section B. All basis set algorithms were run both with and without the MM3 component; in all cases, the reported “GABA+” values include contributions from the underlying macromolecule signal, either explicitly in cases where the MM3 component was modelled (i.e., GABA + MM3co), or implicitly in cases where it was not.

#### 2.3.2 Quality Control: Modelling

The available quality metrics vary between algorithms; all except Osprey report some form of modelling uncertainty (%SD, %CRLB of metabolite estimates, or % Fit Error for the model), and most report SNR and linewidth of water and/or some metabolite components. As specifics of each algorithm’s SNR and linewidth calculation vary, independently derived values are assessed at the processing stage (R2, section 2.2.1). Adopting rather liberal criteria, individual fits were flagged as having failed if either of the following additional criteria were met; in cases where a given metric was not available, the condition is ignored. Criteria below are designated **R3** and **R4** for subsequent usage.

- R3: %SD, CRLB or FitError for GABA+, Glx_diff_ or tCr_edit_off_ estimate exceeded 50% (per ^33,41^, acknowledging that this strategy must be used with caution ^42^)
- R4: Final, scaled estimate for any target metabolite (GABA+/H_2_O, Glx_diff_/H_2_O, GABA+/tC_redit_off_, Glx_diff_/tCr_edit_off_) differing from the median value by more than 5 times the median absolute deviation (MAD) ^43^ for that algorithm; this was intended to capture any poor fits not flagged by any other criteria.

Visual inspection of data, fit outcomes and residuals was also performed, to confirm that no grossly aberrant outcomes eluded the defined rejection criteria. All subsequent analyses are performed after exclusion of individual algorithms’ fits (not entire subject datasets) per these criteria.

### 2.4 Statistical Analysis of Modelling Outcomes

After batch modelling, statistical analysis was performed using locally implemented scripts written in Python (v3.7.3), using the pandas ^44^ (v0.23.3) data analysis framework, with numeric methods from NumPy ^45^, and statistical methods from the SciPy ^46^ (v1.1.0), pingouin ^47^ and statsmodels ^48^ (v0.12.1) libraries.

Scaled estimates for target metabolite, grouped by algorithm, were tested for normality using the Shapiro-Wilk method ^49^, and for comparable distribution of variance between algorithms by the Fligner-Killeen’s test ^50^, both implemented in SciPy. Limits of agreement between pairs of algorithms were derived, along with their 95% confidence intervals, in accordance with the Bland-Altman method ^51^. Estimates grouped by algorithm and manufacturer were compared using Welch’s t-test ^52^, with Holm-Bonferroni correction ^53,54^ for multiple comparisons. An adjusted p-value less than 0.05 was considered significant.

An unconditional linear mixed-effects model was fit to water-referenced GABA+ estimates, using R version 3.5.3 ^55^ with the *lme4* package ^56^ and an implementation derived from ^25^. Vendor, site, algorithm and subject factors were incorporated into the fit, to calculate variance partition coefficients (VPCs) estimating the proportion of total variance attributable to each factor. Significance testing was performed using chi-square likelihood ratio tests, against a null hypothesis simulated by parametric bootstrapping (2000 simulations) ^57^. The Big GABA dataset described herein has previously been assessed with respect to demographics and signal quality ^24^.

For each metabolite of interest, a global median was calculated across all subjects and all algorithms. Subsequently, estimates grouped by site and algorithm were linearly scaled to match the global median, thereby removing broad scaling differences observed between certain sites, vendors and algorithms which would otherwise bias inter-algorithm correlations.

The degree of correlation between MRS-detected GABA estimates and voxel tissue fraction has been shown to be an effective index of GABA estimation accuracy ^35^, given the differing GABA concentrations between grey and white matter ^58–60^. Building on this approach, robust Spearman correlation coefficients between voxel grey matter fraction and GABA+/H_2_O_noTC_estimates were calculated using the ‘skipped’ method ^61,62^, implemented in pingouin, to exclude bivariate outliers. To determine whether correlation coefficients obtained for any one of the algorithms differed significantly from the correlation obtained across all algorithms, z-scored coefficients were compared (two-tailed), with a significance level of p_holm_ < 0.05; similarly, z-scored coefficients were compared between variants without and with the MM3 macromolecule component in the basis set.

Finally, intraclass correlation coefficients (ICCs) were calculated between all algorithms, separately without and with the MM3 component for basis set algorithms, and between pairs of algorithms, using a two-way mixed-effects model for single-rater consistency (ICC(3,1) implemented in pingouin).

## 3 Results

### 3.1 Fit and Residuals

Mean fit outcomes for each algorithm are presented in Figure 2. Note that all basis set algorithms without MM3, except QUEST, show strong residuals in the 3 ppm range; in the case of Osprey (Figure 2f), this appears to have a strong influence on baseline in the vicinity. Variants with MM3 show generally reduced residuals in that range, indicating that the inclusion of a dedicated MM3 basis function leads to more appropriate modelling of the data.

**Figure 2.**
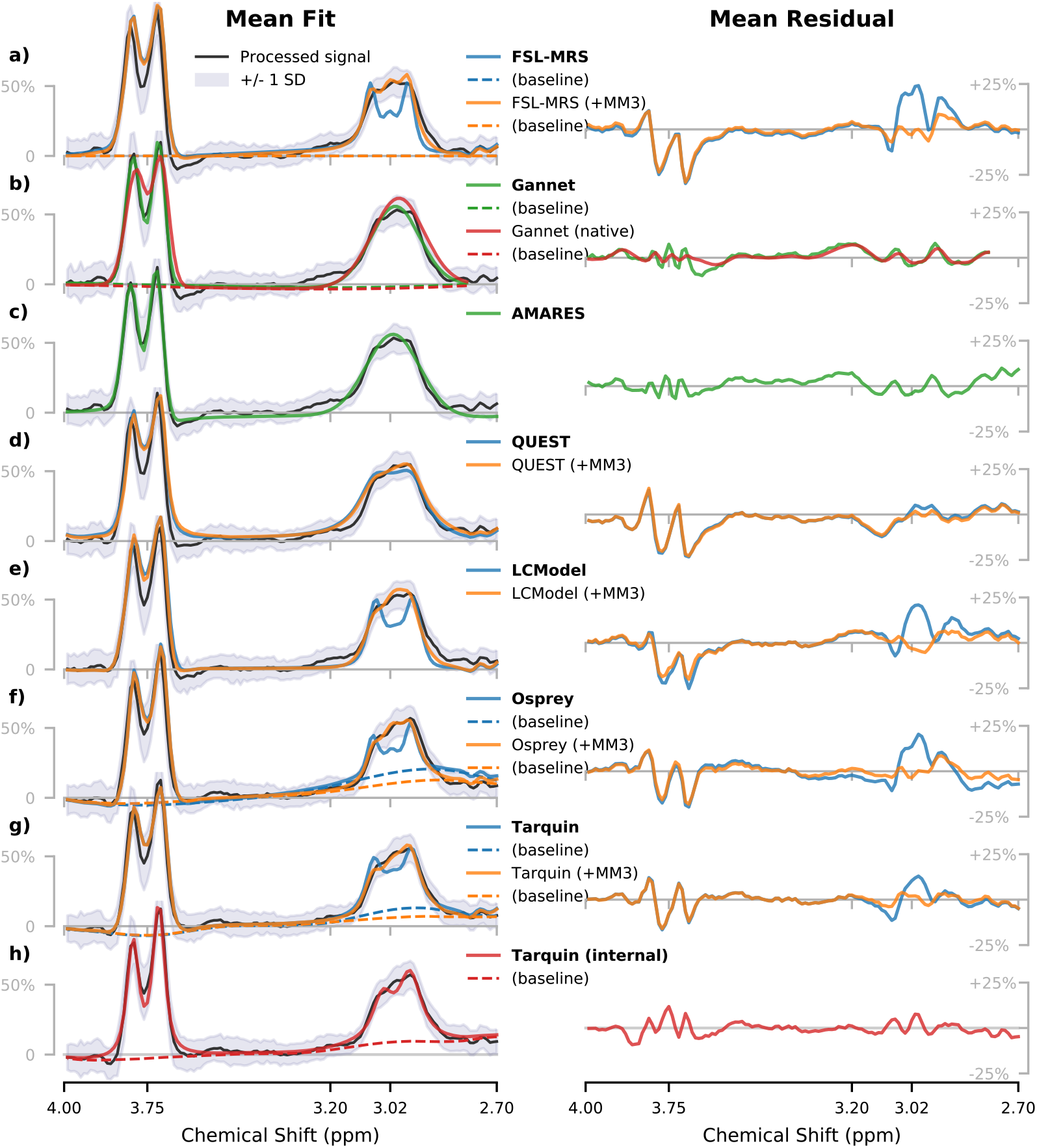
Average metabolite and baseline (where applicable) models with corresponding residuals for the GABA+ edited spectra, for each algorithm. Vertical scaling is normalised; outcomes over the full fit range are presented in Supplementary Figure 8; outcomes split by vendor are presented in Supplementary Figure 9.

A characteristic hump around 3.2 ppm is handled differently by the various algorithms: peak fitting algorithms Gannet and AMARES, (Figure 2b,c) are largely unperturbed, QUEST (Figure 2d) envelopes the entire signal with broader 3.0 ppm peak, while other algorithms fall somewhere in between.

A notable difference between algorithms arises from the differences in baseline estimation practices. While AMARES, QUEST, and LCModel do not include a baseline term in their default settings for MEGA-PRESS and Gannet and FSL-MRS adopt relatively stiff, low-order models, both Tarquin and Osprey attribute a considerable fraction of the edited 3-ppm signal to the baseline. This tendency is mitigated upon the inclusion of the MM3 model.

Finally, there is a distinct pattern to the residuals around the Glx peaks from all basis set algorithms, not present in the fits applying simple peak fitting on a restricted frequency range (Gannet, AMARES).

A summary of basic quality metrics from the fit spectra is presented in Supplementary Figure 6, along with the number of spectra rejected according to the defined criteria (per sections 2.2.1 and 2.3.2).

### 3.2 Statistical Analysis

Shapiro-Wilk testing and subsequent inspection of quantile-quantile (Q-Q) plots indicated that while concentration estimates from most algorithms satisfied the assumption of a normal distribution, several (predominantly Glx_diff_/tCr estimates) deviated slightly from this. Fligner-Killeen tests revealed mismatched variances between several sets of estimates (predominantly relating to QUEST and LCModel +MM3), which motivated the subsequent adoption of Welch’s t-test for groupwise comparisons.

#### 3.2.1 Water-referenced concentration estimates

Comparisons between algorithms for GABA+/H_2_O are summarised in Figure 3, with full details in Supplementary Table 6, and Bland-Altman plots describing limits of agreement in Supplementary Figure 10. The global median estimate for GABA+/H_2_O across all algorithms and subjects was found to be 3.2 ± 0.4 i.u..

**Figure 3:**
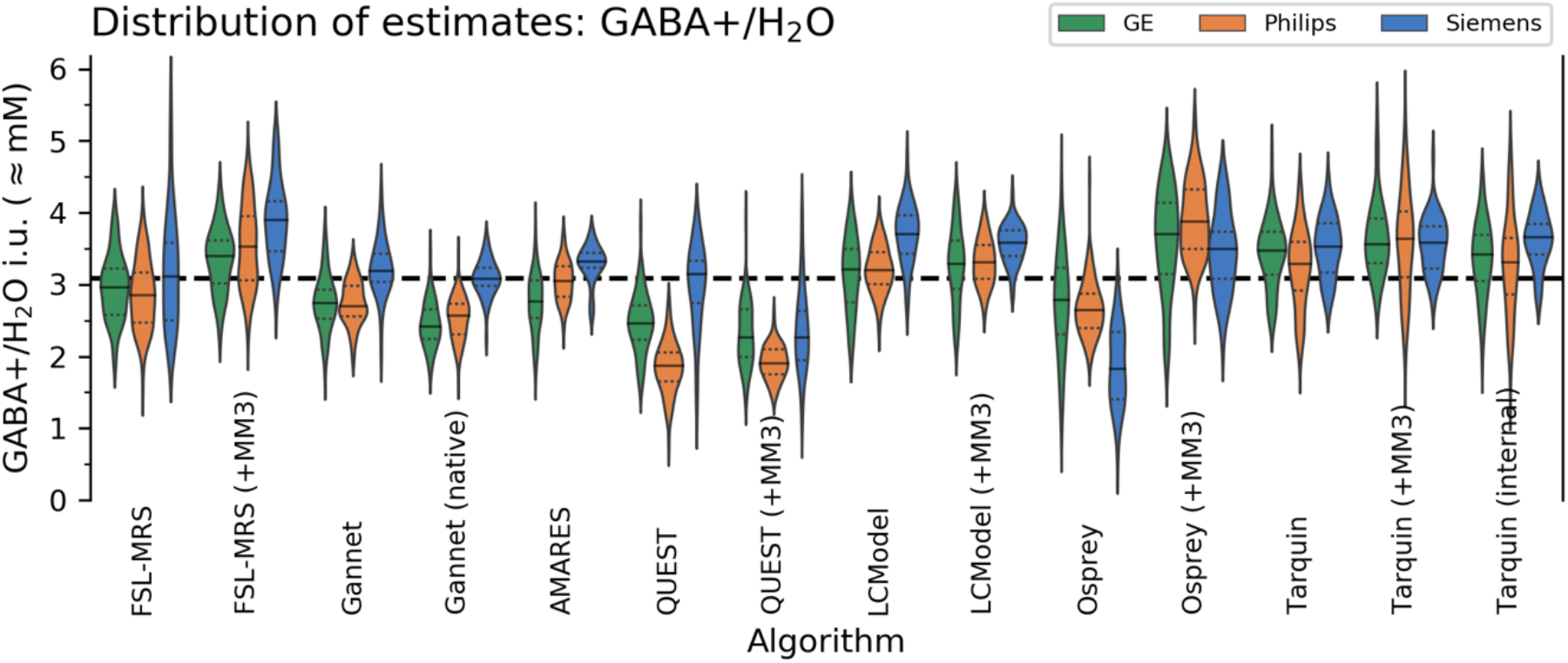
Distribution of GABA+/H_2_O estimates from each algorithm, grouped by manufacturer. Global median is shown in dashed black.

For several quantification algorithms, water-referenced estimates for GABA+/H_2_O were found to be significantly higher from Siemens datasets than from other manufacturers, by a factor of 9-17% (p_holm_<0.001, depending on the algorithm) for FSL-MRS (+MM3), Gannet, Gannet (native), AMARES and LCModel, increased to 41% (p_holm_<0.001) for QUEST. Osprey gave significantly lower estimates for Siemens datasets (−28.0%, p_holm_<0.001). QUEST (and the +MM3 variant) gave significantly lower estimates for Philips datasets (−16.1% and −7.7% respectively, p_holm_<0.001), and AMARES and Gannet (native) gave lower estimates for GE datasets (−9.5%, p_holm_<0.01 and −8.5%, p_holm_<0.05 respectively). Median GABA+ estimate across all algorithms was 5.8% higher for Siemens sites (p_holm_<0.01). All differences are expressed relative to the mean across all subjects for the respective algorithm. No other variants showed significant effects.

Water-referenced Glx_diff_ estimates from all algorithms were significantly higher for Siemens sites: median Glx_diff_/H_2_O across algorithms +15.7%, p_holm_<0.001 relative to group mean. Estimates from Philips sites were somewhat lower (−10.1%, p_holm_<0.01).

For data fit without the explicit MM3 component, the unconditional linear mixed-effects model yielded VPCs of [33.8, 16.4, 6.4, 4.0%] for algorithm, site, subject and vendor factors respectively. In this context, the “subject” factor reflects systematic *within-subject* variation in estimates, while the residual 39.4% accounts for inherent, systematic *between-* subject variation, as well as any other variance which could not be accounted for in the model. Parametric bootstrap testing showed all factors to be significant (p_holm_<0.001).

#### 3.2.2 Metabolite-referenced concentration estimates

Estimates for GABA+/tCr_edit_off_ were consistently higher for GE datasets (+17.3% across algorithms, p_holm_<0.001) and lower for Siemens datasets (−14.3%, p_holm_<0.001). Glx_diff_/tCr_edit_off_ ratios were higher in GE datasets (21.9%, p_holm_<0.001) and slightly lower in Philips (−5.0%, p_holm_<0.05) and Siemens (−6.7%, p_holm_<0.05) datasets. As in section 3.2.1, differences are quoted relative to the mean estimate across all subjects for the respective algorithm. All these trends presented similarly across all modelling algorithms, albeit with varying magnitudes and significance levels.

#### 3.2.3 Grey matter volume fraction correlation

The relationship between estimated GABA+ and grey matter volume fraction is reported in Figure 4, as an index of estimation accuracy. The accuracy of QUEST (without MM3) was found to be significantly below that of other algorithms (p_holm_<0.01), while the QUEST +MM3 variant performed comparably with other algorithms. Otherwise, slight differences observable between algorithms were not statistically significant, and no particular trend is evident between algorithm variants without and with MM3 components.

**Figure 4.**
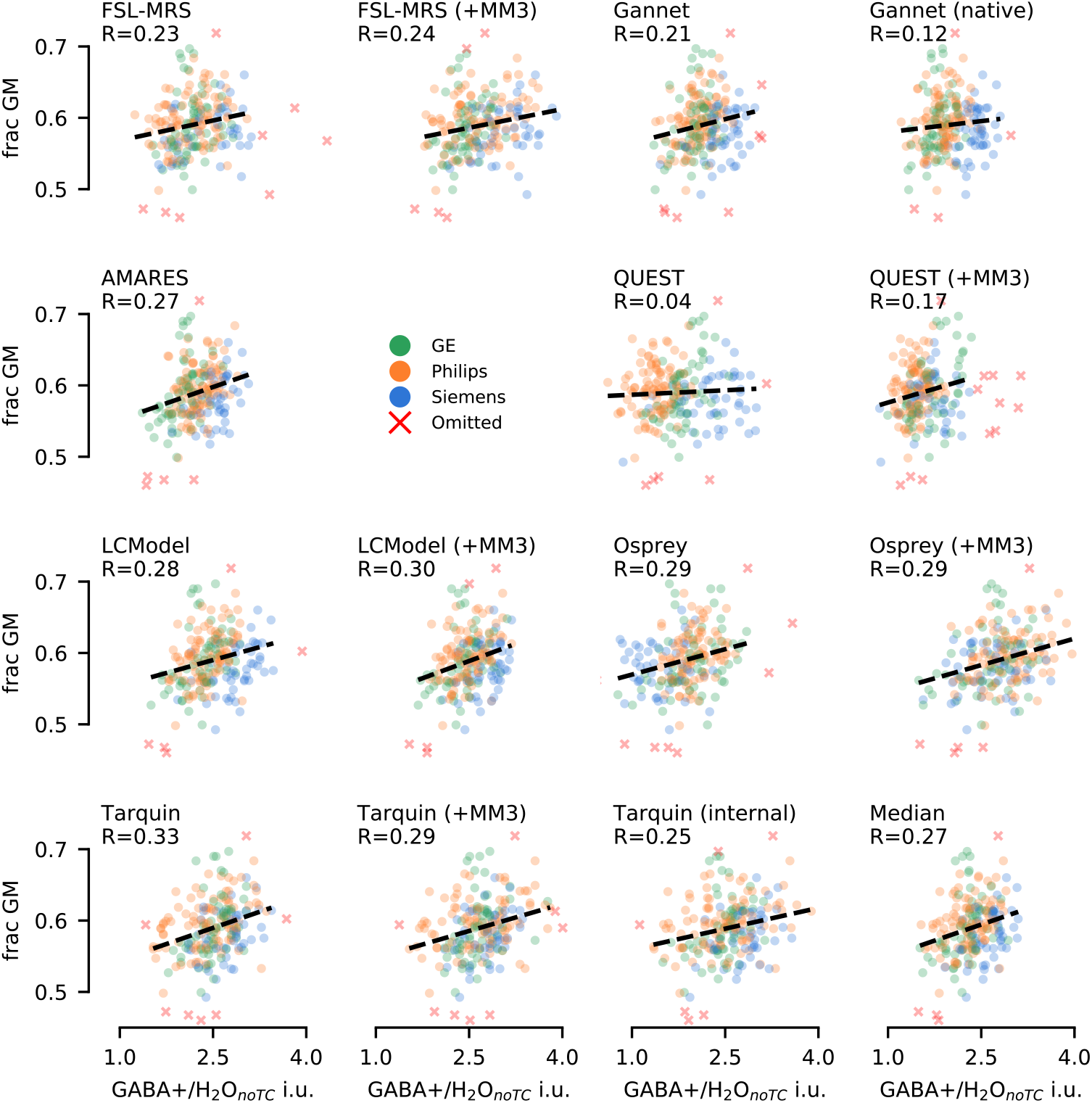
Relationship between GABA+ and grey matter, with different modelling strategies for GABA+. Robust (skipped) correlation coefficients are reported, with line-of-best-fit in dashed black.

#### 3.2.4 Correlational analysis

ICC (single-rater consistency) for GABA+ across all algorithms was 0.38 (95% CI 0.32-0.44) without the MM3 component for basis set algorithms, and increased to 0.44 (95% CI 0.39-0.5) with MM3 included, supporting that the inclusion of this dedicated component is warranted. ICCs between all pairs of algorithms are presented in Figure 5. For fits performed without the MM3 component, GABA+/H_2_O estimates showed moderate correlation between most algorithms (typically on the range r=0.4-0.6; slightly lower when referenced to tCr_edit_off_). Correlations for AMARES, LCModel and Tarquin were significantly stronger (p_holm_<0.01) than the group mean, those for QUEST and Osprey somewhat lower. Inclusion of an MM3 basis set component generally improved concordance with other algorithms for FSL-MRS (p_holm_<0.001) and Osprey, the latter at trend level. However, both time domain basis set algorithms (QUEST and Tarquin) showed reduced concordance (at trend level) upon inclusion of the MM3 component.

**Figure 5.**
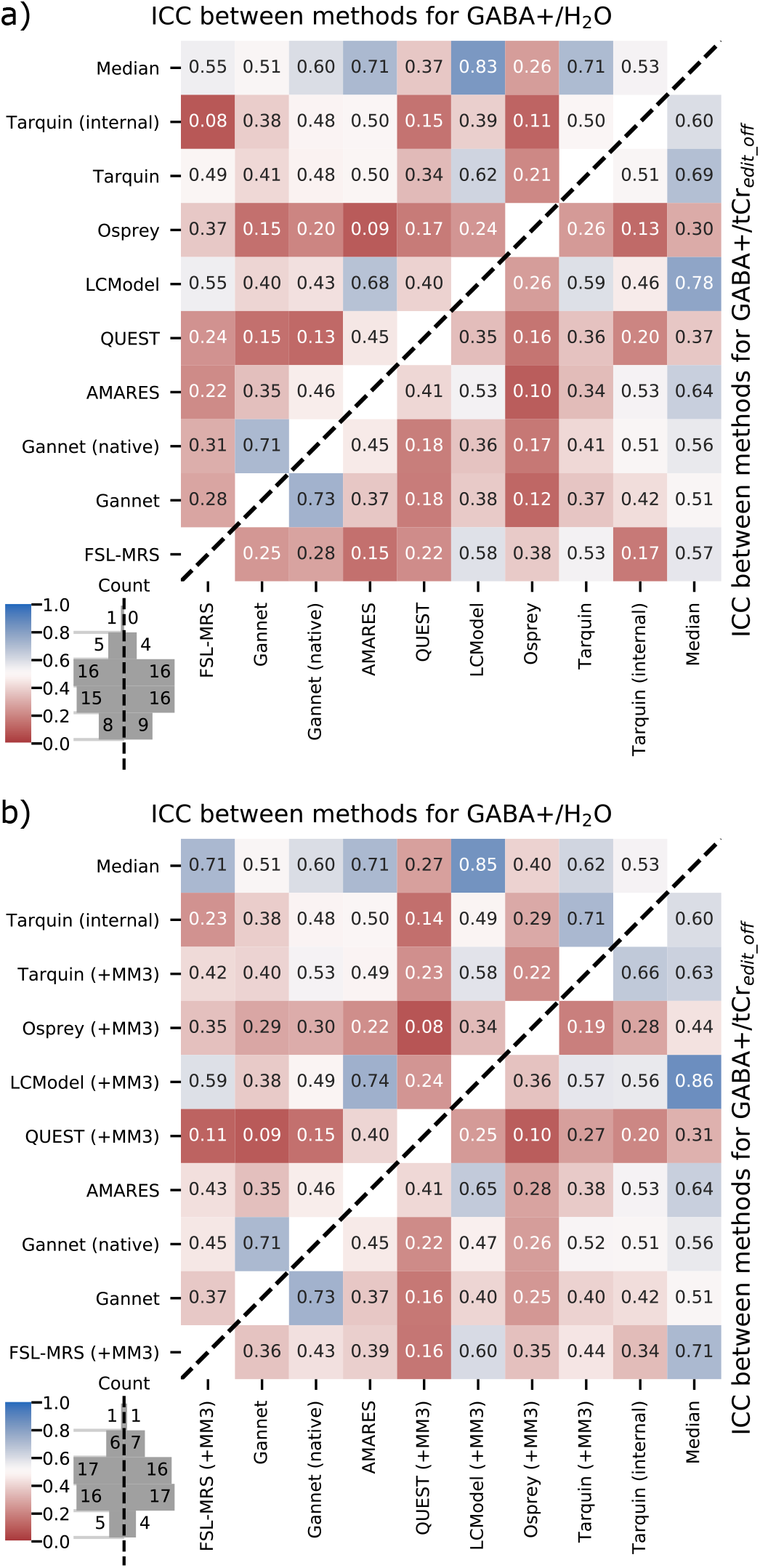
Intraclass correlation coefficients between algorithms, scaled to water (upper left triangle) and tCr_edit_off_ (lower right triangle), with basis set algorithms excluding (a) and including (b) a component representing co-edited macromolecule contribution. “Median” data denotes correlation with the median estimate across all algorithms.

ICCs for additional metabolites and ratios are presented in Supplementary Figure 12; Glx_diff_/H_2_O estimates from the edited spectrum correlated more strongly between algorithms (typically on the range r=0.6-0.8, slightly lower when referenced to tCr_edit_off_).

## 4 Discussion and Conclusions

### 4.1 Quality Control

Basic signal quality metrics (such as SNR and linewidth) and reliability-of-fit estimates (%CRLB, %Fit Error) are often used as the basis for rejecting poor fits. However, as seen in Supplementary Figure 6, these are often not sufficient. Whilst four datasets were deemed to have failed at the processing stage (R1), yielding output barely recognisable as GABA-edited difference spectra, all algorithms “successfully” fit some of these (Supplementary Figure 7), with quality metrics that satisfied all other criteria. We therefore repeat the observation that simply filtering results based on these basic signal metrics is inadequate as a means of quality control; the metrics themselves may have limited reliability, particularly in cases where the model does not accurately reflect the experimental data ^63^. Consideration must also be given to the shape of the data, fit and residuals themselves – either by algorithmic assessment or, if feasible, visual inspection.

### 4.2 Scale Differences between Manufacturers

Previous studies, including ^64^, have explored systematic differences in reported GABA+ estimates between different manufacturers, and their relation to GABA editing efficiency and the contribution of co-edited macromolecules to the measured signal. Furthermore, a previous examination of water-scaled GABA+ estimates on a superset of data also incorporating the present study’s subjects ^24^ identified systematically higher GABA+ estimates from datasets acquired on the Siemens platform, by approximately 29%, which could not be explained in terms of editing efficiency or macromolecule contribution. Observations in the present analysis corroborate this, with most algorithms yielding higher GABA+/H_2_O estimates from Siemens datasets, and all algorithms yielding higher estimates for both Glx_diff_/H_2_O and tCr_edit_off_/H_2_O for Siemens datasets. The fact that this trend is seen across different metabolites and within the edit-OFF sub-spectra gives support to the notion that water reference data from the Siemens implementation may not be optimal for scaling purposes, although it cannot be ruled out that both GE and Philips share a similar mis-scaling.

### 4.3 Macromolecule Fitting

The observed signal around 3 ppm includes substantial contamination (around 50% ^65^) from co-edited signals, including homocarnosine ^66^ with coupling between 1.89 and 3.00 ppm spins, and poorly-characterised components tentatively attributed to lysine-containing macromolecules ^6,67,68^, with coupling between 1.71 and 3.01 ppm spins. This may give rise to large residuals or biased baseline estimates if not considered in the model. Although their impact has been studied by some ^38–40,69,70^, there is currently no consensus on how these should be handled.

The default LCModel configuration (flat baseline) performs surprisingly well by simply ignoring such signals, giving rise to characteristic peaks in the residuals. Meanwhile, the case of Osprey (without MM3co component, Figure 2f) exemplifies the potential baseline distortions and correspondingly reduced GABA+ estimates resulting from these residuals. The result of our ICC analysis across all algorithms suggests that more consistent GABA+ estimates may be obtained by explicitly parametrizing the MM3 contribution in the model. However, using correlation between GABA+ estimates and grey matter fraction as a benchmark, incorporation of the MM3 component did not significantly impact the effectiveness of individual algorithms in measuring differences in GABA+ levels. Moreover, while the supplementary analysis (supplementary section B) suggested improved effectiveness for LCModel after incorporation of soft constraints on MM3 amplitude (whether to GABA or MM0.9), similar performance for this algorithm was obtained by simply modelling a stiff but non-zero baseline, allowing this to absorb some of the MM3 signal. This configuration was also found to be effective for modelling GABA alone, consistent with previously reported findings ^69^ wherein modelling a more flexible baseline in LCModel to selectively remove a portion of the MM3 contribution allowed for closer measurement of GABA rather than GABA+. Estimates from Osprey in similar configurations were comparable, and it has been shown that higher degrees of baseline flexibility cause greater fractions of the 3-ppm signal to be absorbed into the baseline ^40^.

Peak-fitting algorithms (such as Gannet and AMARES) may circumvent this issue somewhat by considering the entire 3.0 ppm GABA+MM signal with a single broad Gaussian model as examined herein; this approach performed comparably with more elaborate models. Furthermore, Tarquin with its internally simulated basis set has two separate Gaussian components representing the GABA+ signal, which are seen to shift and broaden to conform to the shape of the observed GABA+MM signal (Figure 2h). QUEST, similarly, broadened the GABA basis function substantially to more closely envelop the entire GABA+MM signal and (unfortunately) adjacent artefacts (Figure 2d), perhaps accounting for its somewhat lower correlation with grey matter fraction and agreement with other algorithms.

### 4.4 Artefact Rejection around 3.2 ppm

MEGA-edited GABA spectra often exhibit a slight artefactual feature around 3.2 ppm, which can be problematic for fitting algorithms. The origin is uncertain, but potentially related to incomplete subtraction of choline ^71^ or contribution from undetermined other co-edited signals (such as, perhaps, valine-containing macromolecules ^67^ or arginine ^72^). In the present study, the baseline for the Osprey fit (without MM3co) tends to respond to this artefact, inducing a bend in the baseline which appears to cut out a significant part of the real GABA peak (Figure 2f), leading to a likely underestimation of GABA+ area. QUEST appears to broaden the GABA and/or MM3co basis components, incorporating the artefact into the GABA+ estimate and most likely over-estimating the GABA+ signal area (Figure 2d); this effect was most pronounced for Siemens datasets (see Supplementary Figure 9d), where the feature manifests more prominently. FSL-MRS and LCModel both handle the artefact well in the general case, largely rejecting it from both the baseline and metabolite models (Figure 2a,e); this is likely owed to the fact that their default MEGA-PRESS settings prescribe a low-order polynomial baseline (FSL-MRS) or no baseline at all (LCModel). Whilst other basis set algorithms end up somewhere in between (with a degree of contamination from the artefact), peak-fitting algorithms AMARES and Gannet both perform well in this area. Indeed, Gannet (Figure 2b) is the only algorithm to explicitly deal with this artefact, down-weighting some residuals in this region. We suggest that comparably rigid baseline estimation as well as incorporating a Gaussian basis component around 3.2 ppm, with tight constraints on linewidth, shift and amplitude to avoid inadvertently fitting part of the GABA peak, may yield some benefits in this area for other algorithms. Ultimately, further investigation into the underlying signal, and more complete profiling of the co-edited metabolite and macromolecule signals in the region would be preferable.

### 4.5 Glx

Although quantification of Glx from the difference spectrum has been demonstrated to be reliable given suitable quality constraints ^73^, several researchers have highlighted the relatively low concordance between estimates from short-TE PRESS spectra and GABA-edited difference spectra, with estimates from the edit-OFF sub-spectra often found to agree better with the short-TE PRESS ^74–76^; this is unsurprising given that the Glx signal in the short-TE and edit-OFF sub-spectra are subject to similar underlying uncertainties, including MM background and overlapping GSH and aspartyl signals. In comparing Glx quantification between MEGA-PRESS difference and edit-OFF sub-spectra, recent studies ^76,77^ report a correlation around r=0.8; results in the present study show a more moderate correlation, between r=0.34 and 0.69 depending on algorithm: see Supplementary Figure 13; a linear scaling factor is also observed, consistent with recent findings ^78^. It is notable that agreement between algorithms is higher for co-edited Glx than for GABA+, reflecting the better-defined signal seen in the difference spectrum (Figure 2).

With reference to Supplementary Figure 8, all basis set algorithms showed a distinct structure in the residuals around 3.7 ppm, with the model peaks appearing a little to the right of the peaks observed in the data. This is most likely due to the complicated signal patterns around 2.3 ppm in the edited spectrum (resulting from overlapping signals of GABA and co-edited Glu, Gln and GSH), which interact critically with the 3.75 ppm modelling. It is likely that there is a poorly understood baseline fluctuation arising from co-edited macromolecular signals appearing between 1.5 and 2.5 ppm ^39^, which will bias the correct phase estimation of the 2.25 ppm signals, at the expense of getting the phase of the related 3.75 ppm signals right. The peak-fitting algorithms tested, where modelling around 3.7 ppm is not bound to features in other parts of the spectrum (such as around 2.3 ppm), show much lower residuals in the region. It is possible that basis set fitting on a constrained range would mitigate this effect, at the expense of throwing away useful spectral information and hence detracting from the utility of the basis set approach in general. A model which shares lineshape information between the 2.3 and 3.7 ppm Glx peaks but allows a tightly-constrained frequency shift between them may present a reasonable alternative.

### 4.6 Limitations

The basis set adopted in the present study was simulated with ideal excitation pulses, and therefore may not fully model subtle variations in spectral structure between manufacturers. However, the impact of excitation is likely negligible compared to the impact of refocusing, which is appropriately accounted for in the 2D simulations. Vendor-specific excitation pulses may also contribute to subtly different shape and asymmetry of the 3.0 ppm peak and varying manifestation of the 3.2 ppm feature, which may be observed in Supplementary Figure 9.

Whilst the present analysis examines a variety of commonly used implementations representing a range of modelling strategies for GABA-edited spectroscopy data, we note that several other algorithms and implementations are also available to the MRS community, including AQSES ^79^, INSPECTOR ^80^, KALPANA ^81^, OXSA ^82^, spant ^83^ and Vespa ^84^.

Furthermore, many of the packages examined offer extended functionality which may well lead to improved performance in certain circumstances, but this did not align with our approach of adopting recommended/default configurations for all algorithms. Most significantly, many software packages offer the fine-tuning of several aspects of the modelling process, for instance the baseline parametrization. As further examples, jMRUI QUEST offers flexible baseline modelling strategies; FSL-MRS offers independent shift groups which were not assessed; Osprey can additionally simultaneously optimise difference and sum spectra, potentially benefiting from additional spectral information and improved SNR. Both peak-fitting algorithms examined are flexible in their choice of model, with (for example) dual-Gaussian models for the GABA+ signal readily available. An inevitable consequence of adopting default settings in this analysis is that these might not be optimal for data processed with the standard pipeline adopted herein. All the tools tested are highly configurable and offer expert users many possibilities to tune performance optimally for particular datasets, offering the potential for further invention and protection against establishing a possibly incorrect orthodoxy. Nonetheless, this flexibility comes with the caveat that it may also lead to misuse, selective reporting and inappropriate modelling. Moreover, such variability runs counter to efforts to standardize analysis methodology ^1^. While more research into optimized modelling strategies is needed to improve the comparability and robustness of MEGA-PRESS fitting, this work highlights that the complete and accurate reporting of all decisions made during analysis and modelling is immensely important, going even beyond the recently published minimum reporting standards for MRS ^85^.

The present study examines metabolite estimates for each algorithm, relative to water and creatine references obtained using that same algorithm – this reflects typical usage, but means that variations discussed herein are not necessarily driven purely by the metabolite modelling.

Finally, since the findings documented herein are substantiated entirely by in-vivo data, there is no “ground truth” available by which to directly assess the algorithms’ accuracy. Whilst this limitation has been partially addressed by considering the strength of correlation between GABA+ estimates and grey matter fraction as an index of relative accuracy ^35^, a more direct assessment of accuracy could be achieved in further studies involving carefully prepared phantom or synthetic data ^86^, each approach having its own inherent limitations. In either case, meticulous attention to macromolecule baseline, SNR and line shape would be required to ensure transferability of findings to in-vivo applications.

### 4.7 Key Recommendations

Based on these findings, we recommend the following for future studies:

- When applying basis set modelling approaches, special consideration must be given to the co-edited macromolecular signal underlying the 3.0 ppm GABA peak. While appropriate modelling outcomes may be obtained in some cases by entrusting a carefully tuned baseline to capture the entirety of the signal ^40^, a more generalisable approach is simply to routinely incorporate a simulated basis component to represent this signal.
- Care must be taken to ensure consistent behaviour in the presence of commonly observed artefacts, such as the signal around 3.2 ppm. This artefact could be explicitly incorporated into the model, or mitigated with a rigid baseline model, which is less likely to follow the local signal curvature.
- More generally, when inspecting fit outcomes the behaviour of the baseline (where modelled) demands close attention, to ensure that it does not unduly bias modelling of the GABA peak.
- When assessing data acquired at different sites, systematic differences in scale are to be expected and must be considered, regardless of the algorithm applied.

Additionally, we propose four key areas for further systematic investigation in future studies:

1. Robust methods for the generation of large synthetic datasets for validation are necessary to facilitate direct assessment of modelling accuracy. These synthetic data need to be truly representative of in-vivo data, hence the design of their underlying physical signal models will require great attention to detail. Subsequent interpretation must bear in mind that the outcome is likely to be determined by the degree of similarity between the physical model used to generate the data and the model used to decompose it during linear-combination fitting.
2. Detailed exploration of the co-edited macromolecule profile which underlies typical GABA-edited data is required. Metabolite-nulled edited spectra obtained on different hardware platforms could provide the basis for a more informed parametrization of these signals during modelling, and also contribute to accurate and in-vivo-like representation of synthetic data.
3. The origin of the spectral feature around 3.2 ppm requires further investigation: it needs to be determined whether this is a subtraction artefact or an actual real signal (for example from valine-containing macromolecules or other signals hitherto not routinely included in modelling, such as arginine which has a compatible spin system). Clarification of this feature would allow for more appropriate modelling and parametrization of synthetic data.
4. When considering Glx estimates from the difference spectrum, further investigation into the complex interactions of co-edited Glu, Gln, GSH and possibly other signals around 2.3 ppm, and their impact on the 3.75 ppm Glx modelling, would be beneficial. This focus area will benefit from increased insight into the macromolecular background of the GABA-edited spectrum.

### 4.8 Conclusions

Although the observed consistency across algorithms was generally moderate, with pairwise correlation in some cases concerningly weak, we emphasize that more *consistent* estimates are not necessarily more *accurate* estimates: all the algorithms tested (except for QUEST without MM3) were shown to be comparable in their effectiveness in detecting differences in GABA+ concentration. This does however raise some concerns regarding the comparability of findings between different studies, each of which will typically employ a single modelling algorithm, often with divergent processing and prior knowledge and with significantly smaller sample sizes than tested here. This means that the choice of analysis already adds considerable uncertainty and variability.

Improved standardisation of sequence implementation ^27,87^, and adoption of standardised processing pipelines and prior knowledge (e.g. in the form of publicly available basis set libraries) may reduce sources of variation between studies. However, the interaction of baseline, co-edited macromolecule and metabolite signals, and other artefactual signals remains a critical source of variation between algorithms, and within different configurations of the same algorithm. Better characterisation of these signals would allow for more complete modelling (at the risk of over-fitting). Consensus on optimal (or at least, appropriate) control parameters for the respective algorithms would also be beneficial. It may further be of benefit to conceive a “consensus algorithm” to be implemented across software environments, and used as a shared starting point to refine the algorithmic decision making in future iterations of the algorithm. Meanwhile, careful attention to the behaviour of the model with regards to such signals, and rigorous reporting of the configuration employed, are necessary to facilitate meaningful comparisons between studies.

## 5 Acknowledgements

The authors wish to thank Dr Stephen Provencher for his assistance and constructive feedback on the application of the LCModel algorithm.

Analysis was performed within a project funded by ERC grant #249516, which additionally supports the contributions of ARC, LE, KH. Data used in this analysis were previously collected through an international collaborative study funded under NIH grant R01 EB016089.

WTC is supported by funding from Wellcome Trust and the Royal Society (102584/Z/13/Z). PM thanks India –Australia Strategic Biotechnology Funding (BT/Indo-Aus/10/31/ 2016/PKM). RR was supported by the Australian Research Council (DE210100790). GO and RE acknowledge funding support from K99/R00 AG062230, S10 OD021648, P41 EB031771, P41 EB015909, R01 EB016089, R01 EB023963, R01 EB028259 and R21 HD100869.

Graphical abstract illustrated by Laura Garrison (University of Bergen)

## 6 Declaration of Interest

The authors declare no conflicting interests.

## 7 Data Availability Statement

Scripts used for automation and reporting contained in the present manuscript are publicly available here; further dependencies are described within: https://git.app.uib.no/bergen-fmri/analyzing-big-gaba

Spectra analysed in this manuscript were obtained from the publicly available Big GABA repository on NITRC, https://www.nitrc.org/projects/biggaba

Basis sets used in the primary analysis were obtained from the publicly available Osprey package, https://schorschinho.github.io/osprey

## List of Abbreviations

Cho: *choline*
CI: *confidence interval*
Cr: *creatine*
CRLB: *Cramér-Rao lower bound for uncertainty*
diff: *difference (edited) spectrum*
ECC: *eddy-current correction*
FD: *frequency domain*
FI D: *free induction decay (observed time-domain signal)*
FWHM: *linewidth: full width at half maximum*
GABA: *γ-aminobutyric acid*
GABA+: *total edited signal at 3 ppm; GABA with underlying coedited signals*
Gln: *glutamine*
Glu: *glutamate*
Glx: *combined signal of glutamate + glutamine*
GSH: *glutathione*
H_2_O(_noTC_): *water (noTC: referenced without tissue class correction)*
HSVD: *Hankel singular value decomposition*
i.u.: *institutional units*
ICC: *intra-class correlation coefficient*
LCM: *linear combination modelling*
MAD: *median absolute deviation*
MEGA-PRESS: *Mescher–Garwood point-resolved spectroscopy*
MMx(y): *macromolecule signal aroundx(.y) ppm*
NAA: *N-acetylaspartate*
NAAG: *N-acetylaspartylglutamate*
p_holm_: *Holm-Bonferroni adjusted p-value*
ppm: *parts per million*
Q-Q: *quantile-quantile*
R1-4: *adopted rejection criteria; see methods section*
SD: *standard deviation*
SNR: *signal-to-noise ratio*
tCr: *total creatine (creatine+phosphocreatine)*
TD: *time domain*
VPC: *variance partition coefficients*

## Notes

https://git.app.uib.no/bergen-fmri/analyzing-big-gaba

https://www.nitrc.org/projects/biggaba

### Competing Interest Statement

The authors have declared no competing interest.

