## Supplementary Material for "Comparison of seven modelling algorithms for GABA-edited ^1^H-MRS"


A Acquisition Details ... 3
B Exploratory Analysis: co-edited macromolecule handling and baseline interaction ... 4
B.1 Modelling Outcomes ... 6
B.2 Discussion ... 13
C Difference spectrum modelling ... 15
C.1 FSL-MRS ... 15
C.2 Gannet ... 16
C.3 jMRUI: AMARES ... 16
C.4 jMRUI: QUEST ... 17
C.5 LCModel ... 18
C.6 Osprey ... 19
C.7 Tarquin ... 19
D Water Reference: Modelling and Quantification ... 21
D.1 FSL-MRS ... 24
D.2 Gannet ... 24
D.3 jMRUI: AMARES ... 25
D.4 jMRUI: QUEST ... 25
D.5 LCModel ... 25
D.6 Osprey ... 26
D.7 Tarquin ... 27
E Creatine Reference: Edit-OFF sub-spectrum modelling ... 28
E.1 FSL-MRS ... 28
E.2 Gannet ... 29
E.3 jMRUI: AMARES ... 29
E.4 jMRUI: QUEST ... 29
E.5 LCModel ... 29
E.6 Osprey ... 30
E.7 Tarquin ... 30
F Quality Control ... 31
G Fitting Outcomes per algorithm ... 33
H Agreement between algorithms ... 35

I Additional Correlation Analyses ........ 42
I.1 $Glx_{diff}$, $Glx_{edit\_off}$ ........ 43

### A Acquisition Details

| Site | N | Age (years±SD) | Sex F / M | Scanner vendor and model | Software release | Tx/Rx hardware | MEGA-PRESS sequence variant | Phase cycling | Editing inter-leaving | B0 shimming approach | Water suppr. | Spectral width (Hz) | Data points |
|---|---|---|---|---|---|---|---|---|---|---|---|---|---|
| G1 | 7 | 22.9±3.7 | 4 / 3 | GE Discovery MR750w | DV25 | Body coil/32-ch head coil | Interleaved sequence | 2 | 2 | Double-echo GRE | CHESS | 5000 | 4096 |
| G4 | 12 | 25.6±4.5 | 6 / 6 | GE Discovery MR750 | DV25 | Body coil/8-ch head coil | ATSM patch | 8 | 1 | Double-echo GRE | CHESS | 5000 | 4096 |
| G5 | 12 | 25.5±3.7 | 5 / 7 | GE Discovery MR750 | DV25 | Body coil/32-ch head coil | ATSM patch | 8 | 1 | Double-echo GRE | CHESS | 2000 | 2048 |
| G6 | 12 | 24.3±4.2 | 6 / 6 | GE Signa HDx | HD16 | Body coil/8-ch head coil | ATSM patch | 2 | 2 | Double-echo GRE | CHESS | 2000 | 2048 |
| G7 | 12 | 28.1±4.0 | 6 / 6 | GE Discovery MR750 | DV24 | Body coil/8-ch head coil | ATSM patch | 8 | 1 | Double-echo GRE | CHESS | 2000 | 2048 |
| G8 | 12 | 29.7±2.1 | 6 / 6 | GE Discovery MR750 | DV24 | Body coil/8-ch head coil | ATSM patch | 8 | 1 | Double-echo GRE | CHESS | 2000 | 2048 |
| **All G** | **67** | **26.2±4.3** | **33 / 34** | | | | | | | | | | |
| P1 | 9 | 25.0±3.7 | 4 / 5 | Philips Achieva | R5.1.7 | Body coil/32-ch head coil | JHU patch | 16 | 1 | PB-auto | VAPOR | 2000 | 2048 |
| P3 | 12 | 25.1±2.9 | 6 / 6 | Philips Achieva | R3.2.2 | Body coil/32-ch head coil | JHU patch | 16 | 1 | PB-auto | VAPOR | 2000 | 2048 |
| P4 | 12 | 29.2±3.1 | 5 / 7 | Philips Ingenia CX | R5.1.7 | Body coil/32-ch head coil | JHU patch | 16 | 1 | PB-auto | MOIST | 2000 | 2048 |
| P5 | 12 | 24.9±4.3 | 7 / 5 | Philips Achieva TX | R5.1.7 | Body coil/32-ch head coil | JHU patch | 16 | 1 | PB-auto | MOIST | 2000 | 2048 |
| P6 | 8 | 23.1±2.4 | 3 / 5 | Philips Achieva | R3.2.3 | Body coil/8-ch head coil | JHU patch | 16 | 1 | PB-auto | MOIST | 2000 | 2048 |
| P7 | 12 | 27.3±3.7 | 7 / 5 | Philips Ingenia | R5.1.8 | Body coil/32-ch head coil | JHU patch | 16 | 1 | PB-auto | VAPOR | 2000 | 2048 |
| P8 | 12 | 23.6±3.7 | 6 / 6 | Philips Ingenia CX | R5.1.8 | Body coil/32-ch head coil | JHU patch | 16 | 1 | PB-auto | MOIST | 2000 | 2048 |
| P9 | 12 | 23.2±2.0 | 5 / 7 | Philips Achieva | R5.1.7 | Body coil/32-ch head coil | JHU patch | 16 | 1 | PB-auto | VAPOR | 2000 | 2048 |
| P10 | 12 | 25.8±4.6 | 6 / 6 | Philips Ingenia | R5.1.9 | Body coil/15-ch head coil | JHU patch | 16 | | PB-auto | MOIST | 2000 | 2048 |
| **All P** | **101** | **25.4±3.9** | **49 / 52** | | | | | | | | | | |
| S1 | 12 | 25.7±3.7 | 6 / 6 | Siemens Trio | VB17 | Body coil/32-ch head coil | WIP (529) | 16 | 1 | 3D-DESS + manual | CHESS | 4000 | 4096 |
| S3 | 12 | 31.6±3.4 | 9 / 3 | Siemens Prisma | VD13 | Body coil/20-ch head/neck coil | WIP (859D) | 16 | 1 | FAST(EST) MAP | WET | 4000 | 4096 |
| S5 | 12 | 26.5±3.7 | 6 / 6 | Siemens Trio | VB17 | Body coil/12-ch head coil | WIP (529) | 16 | 1 | 3D-DESS | CHESS | 4000 | 4096 |
| S6 | 6 | 26.2±2.0 | 1 / 5 | Siemens Trio | VB17 | Body coil/32-ch head coil | WIP (529) | 16 | 1 | FAST(EST) MAP | WET | 4000 | 4096 |
| S8 | 12 | 24.0±3.5 | 11 / 1 | Siemens Prisma | VE11 | Body coil/64-ch head coil | WIP (859G) | 16 | | 3D-DESS | WET | 4000 | 4096 |
| **All S** | **54** | **26.9±4.3** | **33 / 21** | | | | | | | | | | |
| **Total** | **222** | **26.0±4.1** | **115 / 107** | | | | | | | | | | |

*Supplementary Table 1: Basic demographics, hardware and software parameters for the constituent datasets*

### B Exploratory Analysis: co-edited macromolecule handling and baseline interaction

The incompletely characterised co-edited signals underlying the GABA peak at 3 ppm present a challenge for accurate modelling of the GABA signal in the area. Whilst the main analysis considers the impact of including a simulated basis component representing that macromolecule signal (MM3co: 14 Hz FWHM gaussian centred at 3 ppm), both Osprey and LCModel have the possibility to constrain this component and to adjust baseline flexibility, potentially improving fitting outcomes further. Therefore, a series of variations were assayed as follows:

- **LCModel (no BL):** standard basis set consisting of GABA, Glu, Gln, GSH, NAA, NAAG, MM 0.9, but no MM3co; this is denoted “LCModel” in the main article.
- **LCModel (no BL) +MM3**: as above, with the addition of MM3co to the basis set; this is denoted “LCModel (+MM3)” in the main article
- **LCModel 0.6 BL**: standard basis set, no MM3co component, with a relatively stiff but non-zero cubic spline baseline model:
    - `NOBASE=F` to re-enable baseline modelling (which is disabled by default for MEGA-PRESS when using `SPTYPE=mega-press-3`),
    - `DKNTMN=0.6` to set the baseline knot spacing to 0.6 ppm (the default value is 0.15).
- **LCModel 0.6 BL +MM3**: as above, with the addition of MM3co to the basis set (no explicit constraint)
- **LCModel 0.6 BL 1:1 GABA**: as above, with the addition of a 1:1 soft constraint between GABA and MM3co
    - `NRATIO=2` Two soft constraints will be specified (see below…)
    - `CHRATO(1)=’NAAG/NAA = 0.14 +- 0.15’` represents LCModel’s default soft constraint for NAAG/NAA. LCModel defines several other default constraints,

not applied due to their dependence on components not included in the standard basis set.

- `CHRATO(2)='GABA/MM3co = 1.0 +- 0.1'` applies a soft constraint on the ratio of GABA to MM3co, with an expected value of 1.0 and standard deviation of 0.1.

- **LCModel 0.6 BL 3:2 MM09**: as above, but with a 3:2 soft constraint between MM3co and MM0.9 (and no constraint between MM3co and GABA), as proposed in [1]:
  - `CHRATO(2)='MM3co/MM09ex = 0.66 +- 0.2'`
- **Osprey 0.4 BL**: standard basis set, without the simulated MM3co component. Default 0.4ppm baseline knot spacing. Denoted "Osprey" in the main article.
- **Osprey 0.6 BL**: as above, but with broader 0.6ppm baseline knot spacing
  - `bLineKnotSpace = 0.6`
- **Osprey 0.4 BL +MM3**: same as Osprey 0.4 BL, with the MM3co component included but no constraint applied:
  - `coMM3 = 'freeGauss14'`
- **Osprey 0.4 BL 1:1 GABA**: as with Osprey 0.4 BL +MM3, with a 1:1 soft constraint between MM3co and GABA amplitudes:
  - `coMM3 = '1to1GABAsoft'`
- **Osprey 0.6 BL 1:1 GABA**: as above, with broader 0.6ppm baseline knot spacing – a comparable configuration to the LCModel 0.6 BL 1:1 GABA case.
- **Osprey 0.4 BL 3:2 MM0.9**: as with Osprey 0.4 BL +MM3, but with a 3:2 soft constraint between the simulated MM3co component and MM0.9 amplitude
  - `coMM3 = '3to2MMsoft'`
- **Osprey 0.6 BL 3:2 MM0.9**: as above, with broader 0.6ppm baseline knot spacing – comparable with LCModel 0.6 BL 3:2 MM09

#### B.1 Modelling Outcomes

On visual inspection (refer Supplementary Figure 1), a broader baseline knot spacing for Osprey yields seemingly more complete coverage of the apparent GABA signal when modelled without an explicit MM3 component. However, the stiffer baseline model for Osprey showed a dip in the 3.0 ppm region when the MM3co component was included, suggesting a possible over-estimation of GABA+ area. Inclusion of MM3 in the model resulted in substantially lower residual signal visible around 3.0 ppm for both Osprey and LCModel. Residuals from LCModel modelling showed few discernible differences between the different soft constraint strategies, while the 3:2 constraint to MM 0.9 for Osprey yielded slightly reduced residuals compared to the 1:1 GABA constraint model around 3.0 – 3.2 ppm.

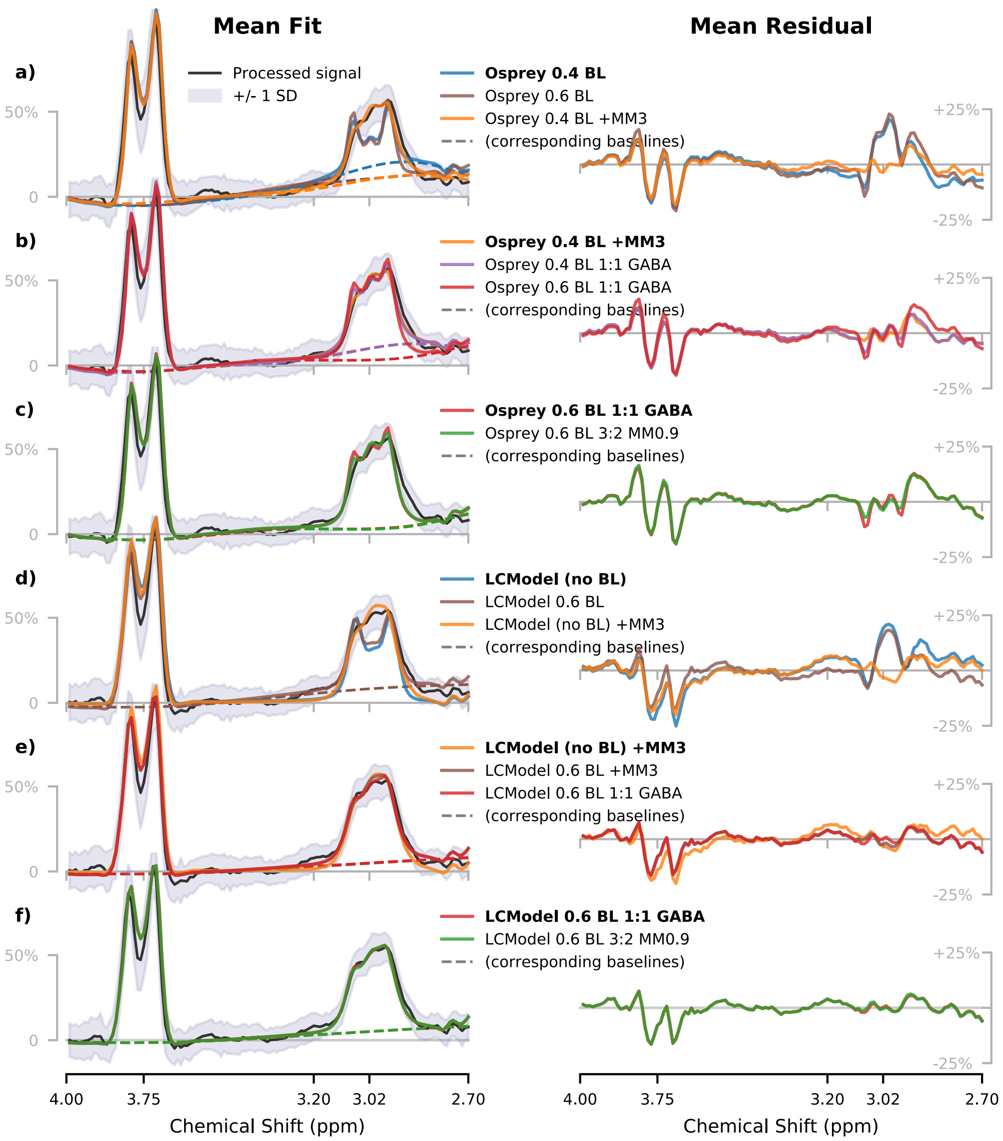


*Supplementary Figure 1: Average metabolite and baseline (where applicable) models with corresponding residuals for each algorithm, baseline model and constraint model in the exploratory analysis. Corresponding fits over the full range are presented in Supplementary Figure 2.*

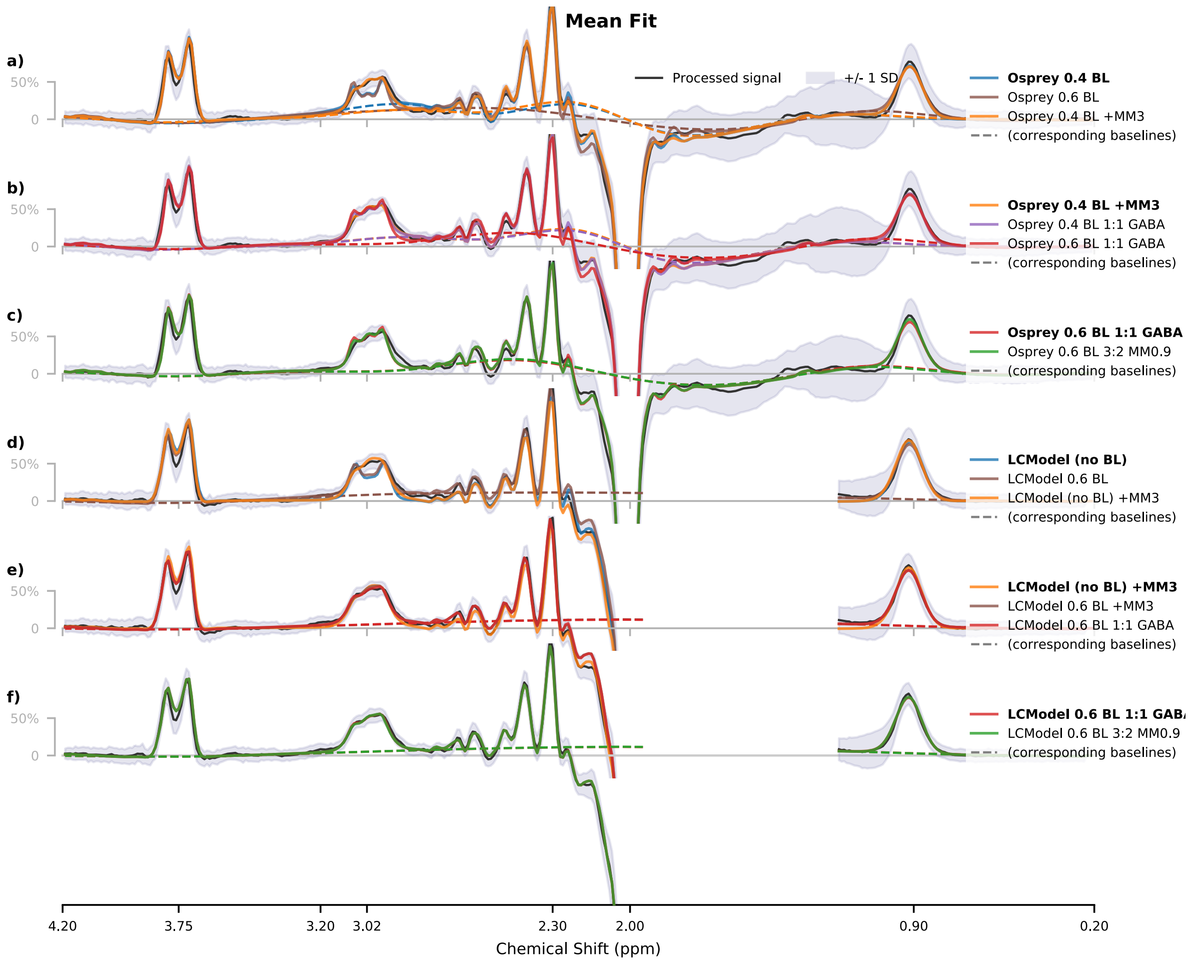


*Supplementary Figure 2: Average metabolite and baseline (where applicable) models for each algorithm, baseline model and constraint model in the exploratory analysis.*

Groupwise statistics are summarised in Supplementary Figure 3 and reported in Supplementary Table 2. Inclusion of the MM3co component appeared to mitigate vendor-specific effects for GABA+/$H_2O$ for both algorithms ($p_{holm} > 0.05$ for most configurations). However, strong, complementary vendor differences were seen for both separate components, GABA/$H_2O$ and MM3co/$H_2O$. For Siemens datasets, GABA/$H_2O$ estimates were systematically lower at trend level (median across configurations -12.6%, n.s.), while MM3co/$H_2O$ estimates were elevated (median across configurations +15.3%, $p_{holm} < 0.01$).

Correlation between GABA+/$H_2O$ estimates and grey matter fraction (Supplementary Figure 4) was comparable across all configurations. While the results suggest somewhat better performance for LCModel 0.6 BL (regardless of the constraint model for MM3), this difference did not approach statistical significance ($p_{holm} >> 0.05$).

For separated $GABA/H_2O$ (Supplementary Figure 5), the LCModel 0.6 BL configuration (having no explicit MM3 component in the model) showed the highest correlation with tissue fraction, although differences between algorithms were not found to be statistically significant. Relative to the $GABA+/H_2O$ estimates from each configuration, only the LCModel 0.6 BL 1:1 GABA configuration (with a 1:1 soft constraint between MM3 and GABA amplitudes) showed significantly degraded correlation ($p_{holm}<0.05$). Differences for Osprey were subtle and non-significant.

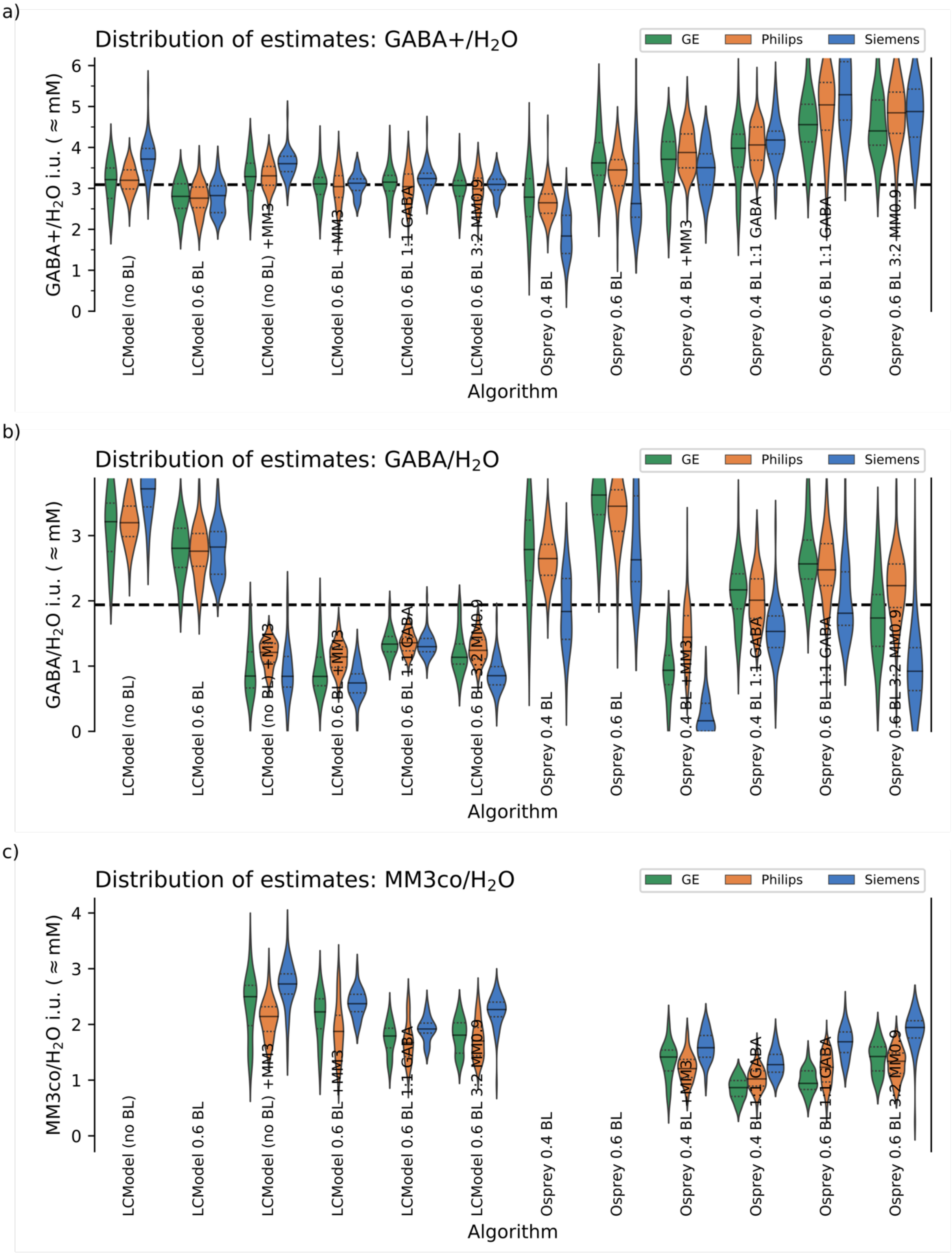


*Supplementary Figure 3: Distribution of metabolite estimates for GABA+ (a), GABA (b), and MM3co (c) obtained from each modelling strategy, grouped by vendor*

| | LCModel (no BL) | LCModel 0.6 BL | LCModel (no BL) +MM3 | LCModel 0.6 BL +MM3 | LCModel 0.6 BL 1:1 GABA | LCModel 0.6 BL 3:2 MM0.9 | Osprey 0.4 BL | Osprey 0.6 BL | Osprey 0.4 BL +MM3 | Osprey 0.4 BL 1:1 GABA | Osprey 0.6 BL 1:1 GABA | Osprey 0.6 BL 3:2 MM0.9 | Median |
|---|---|---|---|---|---|---|---|---|---|---|---|---|---|
| GABA+/H2O mean ±SD | 3.302 ±0.484 | 2.777 ±0.387 | 3.378 ±0.408 | 3.100 ±0.388 | 3.139 ±0.406 | 3.045 ±0.362 | 2.536 ±0.640 | 3.451 ±0.741 | 3.710 ±0.619 | 4.064 ±0.608 | 4.925 ±0.912 | 4.763 ±0.784 | 3.191 ±0.479 |
| GABA+/H2O diff GE | -2.7% | +1.0% | -2.6% | +0.3% | +0.4% | +0.8% | +9.9% | +4.9% | +0.0% | -2.1% | -7.5% | -7.6% | -0.4% |
| GABA+/H2O diff Philips | -3.2% | -0.6% | -2.1% | -1.8% | -3.1% | -1.9% | +4.4% | -0.0% | +4.5% | -0.1% | +2.3% | +1.7% | -2.0% |
| GABA+/H2O diff Siemens | *** +12.5% | +1.7% | ** +6.7% | +0.9% | +3.2% | +1.7% | *** -27.6% | -23.8% | -5.5% | +2.9% | +7.4% | +2.4% | +5.2% |
| GABA/H2O mean ±SD | 3.302 ±0.484 | 2.777 ±0.387 | 1.088 ±0.348 | 0.976 ±0.316 | 1.338 ±0.176 | 1.143 ±0.296 | 2.536 ±0.640 | 3.451 ±0.741 | 1.026 ±0.635 | 1.912 ±0.489 | 2.451 ±0.570 | 1.891 ±0.802 | 1.896 ±0.410 |
| GABA/H2O diff GE | -2.7% | +1.0% | -22.1% | -13.7% | -0.1% | -0.7% | +9.9% | +4.9% | -8.8% | +13.4% | +4.7% | -8.1% | +8.6% |
| GABA/H2O diff Philips | -3.2% | -0.6% | ** +11.2% | *** +16.9% | +1.4% | * +8.6% | +4.4% | -0.0% | *** +40.6% | +5.1% | +1.0% | *** +18.1% | +1.5% |
| GABA/H2O diff Siemens | *** +12.5% | +1.7% | -22.5% | *** -24.0% | -3.0% | *** -25.3% | *** -27.6% | -23.8% | *** -84.2% | *** -19.9% | ** -26.1% | *** -51.3% | -12.6% |
| MM3co/H2O mean ±SD | | | 2.321 ±0.484 | 2.117 ±0.465 | 1.769 ±0.322 | 1.855 ±0.378 | | | 1.358 ±0.320 | 1.031 ±0.269 | 1.225 ±0.366 | 1.444 ±0.379 | 1.746 ±0.411 |
| MM3co/H2O diff GE | | | +7.6% | +5.1% | +1.3% | -2.6% | | | +4.2% | *** -16.0% | *** -23.1% | -1.4% | -0.6% |
| MM3co/H2O diff Philips | | | ** -7.7% | -11.5% | -5.5% | -8.2% | | | * -11.2% | -0.9% | +0.5% | ** -7.2% | -6.4% |
| MM3co/H2O diff Siemens | | | *** +17.5% | *** +12.0% | ** +8.4% | *** +22.2% | | | *** +16.5% | *** +23.9% | *** +37.9% | *** +34.7% | ** +15.3% |

*Supplementary Table 2: Mean concentration estimates (institutional units, ≈mM), for each algorithm in the exploratory analysis. Estimates grouped by vendor are expressed as a % difference relative to the mean across all subjects for the respective algorithm; significance indicated by *, **, *** for $p_{holm}$ < .05, .01 and .001 respectively*

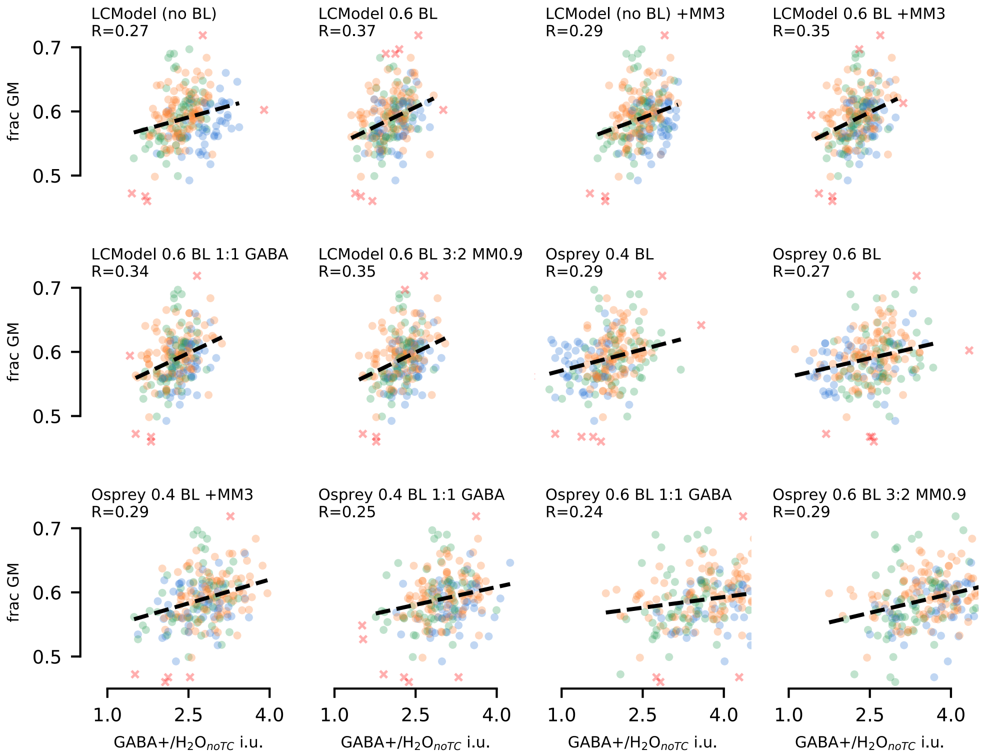


*Supplementary Figure 4: Relationship between GABA+ and grey matter, with different baseline and soft constraint parameters for the MM3 component. Robust (skipped) correlation coefficients are reported, with line-of-best-fit in dashed black*

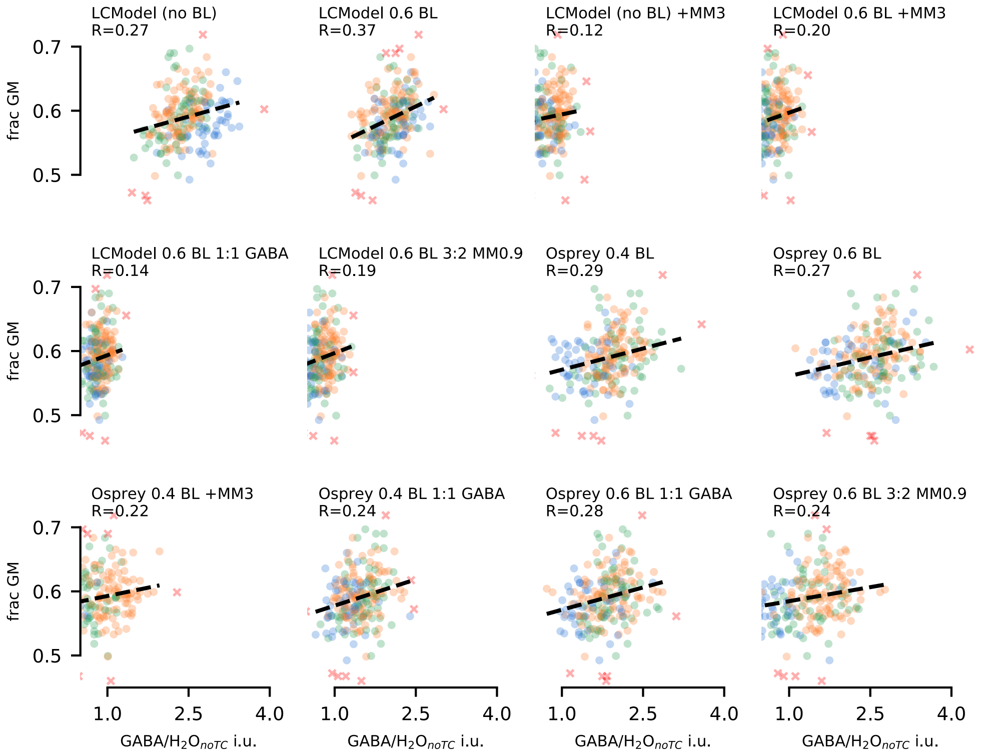


*Supplementary Figure 5: Relationship between GABA and grey matter, with different baseline and soft constraint parameters for the MM3 component. Robust (skipped) correlation coefficients are reported, with line-of-best-fit in dashed black*

#### B.2 Discussion

Whilst a flexible baseline can provide a visually appealing model, i.e., reduce apparent residuals, this does not necessarily lead to more accurate metabolite estimates. This is particularly apparent for the case of Osprey (without the MM3co component), where residuals associated with macromolecule contamination and the 3.2 ppm artefact pull the baseline far into the peak at 3 ppm, potentially leading to a possible under-estimation of the corresponding GABA+ signal. In the present study, stiffer baseline models appeared to be more robust towards small artefacts, allowing the model to reasonably follow the Gaussian contour of the edited GABA+ signal without absorbing too much of it into the baseline.

Nonetheless, using correlation with grey matter fraction as a benchmark, the performance of each algorithm without an explicit MM3 basis component, allowing a stiff but non-zero baseline to absorb the macromolecule signal, was comparable with more elaborate constrained configurations for assessment of GABA+. Moreover, when reporting GABA separately, such a baseline model showed a tendency towards better performance (for LCModel) than other approaches. All configurations except LCModel 0.6 BL 1:1 GABA were able to assess GABA/$H_2O$ separately with comparable effectiveness to GABA+/$H_2O$ from the same algorithm. However, we note that elevated MM3 content in white matter [2,3] may moderate correlation in the case of GABA+. Divergent outcomes for separate GABA/$H_2O$ and MM3co/$H_2O$ estimates across different vendors may be of concern, and requires further attention.

### C Difference spectrum modelling

Further details on specific parameters adopted for modelling the difference spectrum by each algorithm are presented here; this supplements details in the main article, section 2.3.

Basis set construction is described in the main paper, section 2.3.1. Basis sets were exported both in their original resolution (8192 samples at 4 kHz sweep width) and after resampling to match the resolution of the acquired data for each manufacturer/site, to cater for those algorithms which were unable to resample effectively internally (QUEST, Tarquin). The LCModel basis set format was used as an intermediary for import into LCModel, Tarquin and FSL-MRS, while individual jMRUI-format text files were required for assembly into a jMRUI-compatible metabolite list for QUEST.

#### C.1 FSL-MRS

FSL-MRS [4] models data as a linear combination of basis components in the frequency domain, using Bayesian statistics for optimization. Version 1.1.1 was used, which includes updates to the calculation of water-scaled estimates. The standard basis sets as described in section 2.3.1 are used, with the recommended Metropolis-Hastings algorithm and the default fit range (0.2-4.2 ppm).

A single set of shift and line-broadening parameters (for Lorentzian and Gaussian components of a Voigt lineshape model) was applied to all metabolites in the basis set: no additional shift groups were defined.

Default baseline model is a second-order polynomial fit over the defined fit range. To allow FSL-MRS to yield water-scaled estimates, NAA+NAAG was defined as an internal reference for basis set scaling, and fixed tissue content (50% white/50% grey matter, 0% CSF) were supplied. These factors were later reversed (see sections 2.4.1 and D.1).

Typical invocation of FSL-MRS for GABA-edited difference spectra was with the following parameters:

```
--basis ge_diff_8192_4000.basis
--data S12_GABA_68.diff.nii.gz
--h2o S12_GABA_68.ref.nii.gz
```

```
--output S12_GABA_68.diff
--overwrite
--algo MH
--TE 68
--TR 2.0
--tissue_frac .5 .5 0
--internal_ref NAA NAAG
--verbose
--report
--combine NAA NAAG
--combine Glu Gln GSH
--combine GABA MM3co

Defaults:
  --ppmlim:           (0.2, 4.2)
  --baseline_order:   2
  --metab_groups:     0
  --h2o_scale:        1.0
```

#### C.2 Gannet

Gannet [5] models data by fitting peaks in the frequency domain. Version 3.1 was used, with the 'GABA+Glx' model applied to the difference spectrum between 2.79 and 4.1 ppm. This model fits the GABA+ peak with a single Gaussian peak around 3.02 ± 0.05 ppm and Glx as a pair of Gaussian peaks at 3.71 ± 0.02 ppm and 3.79 ± 0.02 ppm, and includes terms to characterize the baseline. The Gannet baseline is modelled as a linear slope with sinusoid and cosine terms, periodic at 2.62 ppm (i.e., $A * (f - f_0) + B * \sin( \pi * f / 1.31 / 4 ) + C * \cos( \pi * f / 1.31 / 4 )$ where $f$ is the frequency (in ppm) and $f_0$ the offset to the first modelled Glx peak (±3.71 ppm).

GannetFit was run on data from the standardised processing pipeline (reported as "Gannet"), and on data processed with Gannet's own default pipeline, denoted "Gannet (native)". The latter applies zero-filling (to 0.062 Hz spectral resolution), and line-broadening (3 Hz), neither of which are performed in the standardised processing pipeline.

#### C.3 jMRUI: AMARES

jMRUI [6,7] is a Java-based package for time domain analysis of MRS and MRSI data, implementing several modelling algorithms. For the present study, jMRUI v6.0 beta was used. Due to compatibility issues with contemporary operating systems, all jMRUI operations were performed on a dedicated 32-bit Debian Linux virtual machine.

AMARES [8] was configured with a GABA+Glx model equivalent to that of Gannet for fitting of the difference spectra, with additional components for NAA and the Glx C4 spins. This incorporated a simple Gaussian model for GABA+ (3.02 ± 0.05 ppm), and a dual-gaussian model for Glx (3.71 ± 0.02 ppm, 3.79 ± 0.02 ppm) – note the close similarity to the Gannet model. The model also used Lorentzian signals to fit the negative NAA (2.0 ± 0.02 ppm) and the Glx C4 spins (2.29 ± 0.02ppm, 2.39 ± 0.02 ppm). Soft constraints were applied on linewidth (5-15 Hz for all Glx, 5-25 Hz for GABA, 5-20 Hz for NAA) and relative phase (±10 degrees, offset by 180 degrees in the case of NAA). No constraints were applied between individual components. The Glx C4 components were included for completeness of modelling, but did not contribute to the reported Glx estimates. Since input data had already been phase corrected, global zero-th and first-order phase were set to zero. Baseline modelling was not performed.

##### C.4 jMRUI: QUEST

QUEST [9,10] is a time domain linear-combination modelling algorithm, implemented within jMRUI. Standard basis set components for each manufacturer and spectral resolution, as described in section 2.3.1, were imported in jMRUI text format and manually aligned for NAA @ 2.0 ppm, before assembly into a jMRUI-format metabolite list.

In earlier testing with the supplied NMRScope-B tool and the supplied megapress_2edit_voi_1pws protocol (configured for TE = 68 ms, TE1 = 13.1 ms, water suppression at 4.7 ppm and 15 ms, 180º gaussian editing pulses at 1.9 and 7.5 ppm adjusted to an “observation offset” of 2 ppm), both the NAAG and GSH components consistently failed to simulate: a complete basis set could not be produced with this tool alone.

The assembled metabolite lists were subsequently used to model batches of data grouped by manufacturer and spectral resolution. The QUEST algorithm was invoked with the zero-order phase fixed to 0 degrees. Other parameters were according to default: no first-order phase; damping factor $\alpha$ (proportional to linewidth, $\Delta$FWHM = $\alpha/\pi$) and frequency shift, per basis function, set on the range -10 to 40 Hz and -10 to 10 Hz, respectively; no baseline terms were included.

#### C.5 LCModel

LCModel [11] models data with a linear combination of basis sets. Version 6.3-1P was used, with the standard basis set described in section 2.3.1. An initial fit was performed according to recommendations in the LCModel manual, with the "special type" set for MEGA-PRESS, on the range 0.2-4.2 ppm with a gap from 1.2-1.95 ppm; water scaling enabled, ECC disabled. This mode of operation assumes a completely flat baseline. Outcomes for this case are labelled as "LCModel". For comparability, the analysis including a basis component representing macromolecule contributions around 3 ppm ("LCModel (+MM3)") leaves all other factors unchanged, including the flat baseline assumption. Additional analyses incorporating a stiff but non-zero baseline and soft constraint models on the MM3 component are considered in a supplementary analysis, section B.

LCModel was invoked with the following parameters for modelling GABA-edited difference spectra:

- `SPTYPE='mega-press-3'` sets the "special type" appropriately for MEGA-PRESS. This implies a flat baseline (`NOBASE=T`), the use of NAA @ 2.01 ppm as a reference for basis set scaling (`wsmet='NAA', wsppm=2.01`), and default correction factor for attenuation of NMR-visible water, ATTH2O = 0.43 (suitable for TE = 68 ms).
- `PPMST=4.2, PPMEND=0.2` model on the range 0.2-0.4 ppm…
- `PPMGAP(1,1)=1.95, PPMGAP(2,1)=1.2` …but exclude the 1.2-1.95 ppm range from modelling
- `DOWS=T` do water scaling
- `DOECC=F` disable ECC

#### C.6 Osprey

Osprey [12] implements a frequency domain basis set fit; version 1.0.1.1 was used, with default fit and reconstruction settings: separate fit on the range 0.2-4.2 ppm, incorporating macromolecule and lipid components in the basis set but without adding a co-edited MM3 peak. Knot spacing for the spline baseline was set at 0.4 ppm. Outcomes according to these settings are labelled "Osprey". The +MM3 variation (Osprey (+MM3)) includes the MM3co basis set component without additional constraints on amplitude. Further variations with respect to baseline knot spacing and soft constraints on MM3 amplitude are considered in a supplementary analysis, section B.

#### C.7 Tarquin

Tarquin [13,14] is a basis set approach which extends the QUEST algorithm for estimation of amplitudes in the time domain. A local build of Tarquin v4.3.11 was used, with the standard basis set per section 2.3.1.

Processed data and basis sets were supplied to Tarquin in the corresponding LCModel file formats; a minor bugfix (subsequently published upstream) was required for LCM basis set import (https://github.com/martin3141/tarquin/commit/cdc44df).

NAA was used as a reference for basis set scaling. Appropriate TE, sweep width and centre frequency (`echo`, `fs`, `ft`) were specified on the command line. Automatic phasing and referencing (`auto_phase` and `auto_ref`) were disabled. The start point for analysis of the difference spectra (`start_pnt`) was set at 5 ms (calculated as 0.005 times the sample frequency). Eddy-current correction was disabled (`--water_eddy false`). HSVD was performed (per defaults) to remove residual water. The visual baseline (Figure 2f) is modelled from the residuals, smoothed with a cosine kernel. Outcomes from this quantification are reported as "Tarquin".

Tarquin can internally simulate basis sets on demand, using an implementation of the density matrix formulation of NMR [15,16] and the parameters of [17]; an additional analysis was performed using this internally generated basis set: `--int_basis megapress_gaba` for the difference spectra. This is simulated assuming a simplified system of uncoupled spins, similar to that used by peak-integration algorithms, comprising GABA_A (3.04 ppm), GABA_B (2.95

ppm), GLX_A (2.299 ppm), GLX_B (2.400 ppm), GLX_C (3.707 ppm), GLX_D (3.789 ppm), NAA (2.0 ppm). Outcomes for this fit are reported as “Tarquin (internal)”.

Typical parameters for tarquin, modelling GABA-edited difference spectra:

```
tarquin
--format lcm
--input S12_GABA_68.diff.RAW
--input_w S12_GABA_68.diff.REF

--echo 0.068
--fs 5000
--ft 127714400
--start_pnt 25
--pul_seq mega_press

--auto_phase false
--auto_ref false
--water_eddy false

--w_att 0.76
--w_conc 35880

--ext_pdf true
--output_txt S12_GABA_68.diff.tarquin.txt
--output_csv S12_GABA_68.diff.tarquin.csv
--output_pdf S12_GABA_68.diff.pdf
--output_fit S12_GABA_68.tarquin.fit
```

For “Tarquin” (with standard basis set, with or without MM3):

```
--basis_lcm ge_diff_4096_5000.basis
```

Or for “Tarquin (internal)”:

```
--int_basis megapress_gaba
```

### D Water Reference: Modelling and Quantification

While all algorithms yielded estimates scaled to water, specifics of the adjustments and correction factors available varied considerably between algorithms; correction according to tissue class was only natively available in a few algorithms (Gannet, Osprey, FSL-MRS). To ensure a fair comparison, scaling as documented for the respective algorithms was first reversed to yield a raw ratio of metabolite area over water area, adjusted by integer scaling factors to account for differing conventions during sub-spectral combination. Subsequently, water-scaled estimates in pseudo-absolute molar units, accounting for differing properties of water in each segmented tissue class, were evaluated according to equation 14 of [18]. Note that additional terms proposed in that paper to account for different metabolite signal relaxation rates in GM and WM were not included, due to currently limited literature on per-compartment relaxation properties of GABA. Constant scaling terms for assumed editing efficiency and macromolecule contamination were applied, per Equation 1. Water-scaled, tissue-class corrected molar concentration estimates are hereafter denoted "/$H_2O$".

$$[\mathrm{M}]_{/\mathrm{H_2O}} = \frac{S_\mathrm{M}}{S_{\mathrm{H_2O}}} \cdot [\mathrm{H_2O}] \cdot \frac{\#H_{\mathrm{H_2O}}}{\#H_{\mathrm{Met}}} \cdot \frac{\sum_{TC}^{GM,WM,CSF} f_{TC,vol} \cdot \beta_{TC} \cdot \mathrm{R}_{\mathrm{H_2O,TC}}}{\left(1 - f_{CSF,vol}\right) \cdot \mathrm{R}_{\mathrm{Met}}} \cdot \frac{MM}{\kappa}$$

*Equation 1*

Where

$$\mathrm{R}_{\mathrm{H_2O,TC}} = \left[1 - \exp\left(\frac{-\mathrm{TR}}{\mathrm{T1}_{\mathrm{H_2O,TC}}}\right)\right] \cdot \exp\left(\frac{-\mathrm{TE}}{\mathrm{T2}_{\mathrm{H_2O,TC}}}\right)$$

$$\mathrm{R}_{\mathrm{Met}} = \left[1 - \exp\left(\frac{-\mathrm{TR}}{\mathrm{T1}_{\mathrm{Met}}}\right)\right] \cdot \exp\left(\frac{-\mathrm{TE}}{\mathrm{T2}_{\mathrm{Met}}}\right)$$

Full parameters are presented in Supplementary Table 3. Individual tissue fractions were obtained from [19], having been originally derived using the GannetSegment module [20] with segmentation of the corresponding per-subject structural $T_1$-weighted image performed using the unified tissue segmentation algorithm of SPM12 [21].

Concentration estimates scaled to water but with no adjustment for tissue class (assuming pure water concentration per eq(3) of [22]) were also calculated, hereafter denoted "/$H_2O_{noTC}$"; see Equation 2.

$$[\mathrm{M}]_{/\mathrm{H_2O_{noTC}}} = \frac{S_{\mathrm{M}}}{S_{\mathrm{H_2O}}} \cdot [\mathrm{H_2O}] \cdot Vis_{\mathrm{H_2O}} \cdot \frac{\#H_{\mathrm{H_2O}}}{\#H_{\mathrm{Met}}} \cdot \frac{1 - \exp\left(\frac{-\mathrm{TR}}{\mathrm{T1_{H_2O}}}\right)}{1 - \exp\left(\frac{-\mathrm{TR}}{\mathrm{T1_{Met}}}\right)} \cdot \frac{\exp\left(\frac{-\mathrm{TE}}{\mathrm{T2_{H_2O}}}\right)}{\exp\left(\frac{-\mathrm{TE}}{\mathrm{T2_{Met}}}\right)} \cdot \frac{MM}{\kappa}$$

*Equation 2*

Subsequent sections D.1 to D.7 detail the water scaling procedure available within the individual algorithms; scaling factors described therein are reversed before applying Equation 1 and Equation 2 to the raw ratio of intensities.

| Parameter | Value | Source and Remarks |
|---|---|---|
| $S_{\mathrm{M}}$ | | Modelled metabolite signal intensity |
| $f_{TC,vol}$ | | Tissue volume fractions, for TC of [GM, WM, CSF] |
| **Water** | | |
| $S_{\mathrm{H_2O}}$ | | Modelled water signal intensity |
| $[\mathrm{H_2O}]$ | 55510 mM | Molar concentration of pure water |
| $Vis_{\mathrm{H_2O}}$ | 0.65 | Assumed MR-visible water, approximated for pure WM from [23] |
| $\beta_{GM}$ | 0.78 | Tissue-dependent water content for GM [23] |
| $\beta_{WM}$ | 0.65 | Tissue-dependent water content for WM [23] |
| $\beta_{CSF}$ | 0.97 | Tissue-dependent water content for CSF [23] |
| $\mathrm{T1_{H_2O}}$ | 1.1 s | Average of WM and GM from [24] |
| $\mathrm{T1_{H_2O,\,GM}}$ | 1.331 s | [24] |
| $\mathrm{T1_{H_2O,\,WM}}$ | 0.832 s | [24] |
| $\mathrm{T1_{H_2O,\,CSF}}$ | 3.817 s | [25] |
| $\mathrm{T2_{H_2O}}$ | 0.095 s | Average of WM and GM from [24] |
| $\mathrm{T2_{H_2O,\,GM}}$ | 0.11 s | [24] |
| $\mathrm{T2_{H_2O,\,WM}}$ | 0.0792 s | [24] |
| $\mathrm{T2_{H_2O,\,CSF}}$ | 0.503 s | [26] |
| $\#H_{\mathrm{H_2O}}$ | 2 | Number of protons contributing to measured water signal |
| **GABA** | | |
| $\mathrm{T1_{Met}}$ | 1.31 s | [27] |
| $\mathrm{T2_{Met}}$ | 0.088 s | [28]; note that [29] report slightly shorter T1 and T2 for GABA (0.8, 0.13 s respectively). |
| $\#H_{\mathrm{Met}}$ | 2 | Number of protons contributing to measured GABA signal |
| MM | 0.45 | Correction factor for the assumed proportion of macromolecule signal in the measured GABA+ peak [30]; [31] reports 0.46 from simulations. Commonly-assumed value, but note that this factor is expected to be implementation-dependent, based on length and shape of editing pulses (and any applied macromolecule suppression). |
| $\kappa$ | 0.5 | Assumed editing efficiency of GABA [30]; note that [31] reports 0.41 for MEGA-PRESS. |
| **Glx** | | |
| $\mathrm{T1_{Met}}$ | 1.23 s | [32] |
| $\mathrm{T2_{Met}}$ | 0.18 s | [33] |
| $\#H_{\mathrm{Met}}$ | 1 | Gannet and AMARES model one proton around 3.75 ppm; basis set methods incorporate signals from four more protons, around 2.15, 2.45 ppm. |
| MM | 1.0 | |
| $\kappa$ | 0.4 | Gannet adopts this value, citing FID-A simulation at TE=68ms |
| **tCr** | | No MM/$\kappa$ factors for tCr (edit-OFF) |
| $\mathrm{T1_{Met}}$ | 1.350 s | [34] |
| $\mathrm{T2_{Met}}$ | 0.154 s | [34] |
| $\#H_{\mathrm{Met}}$ | 5 | |

*Supplementary Table 3 Parameters for water-scaled metabolite estimates*

#### D.1 FSL-MRS

FSL-MRS models the time-domain water reference data with a Voigt decay function [35], then takes the integral of that function's Fourier transform over the 1.65-7.65 ppm range.

Initial molar concentration estimates were obtained assuming 50% GM, 50% WM (i.e., $f_{GM,vol}$ = 0.5 and $f_{WM,vol}$ =0.5). Molar scaling in FSL-MRS is equivalent to Equation 1, but adopting slightly different relaxation parameters as detailed in Supplementary Table 4.

| Parameter | Value | Source and Remarks |
|---|---|---|
| **Water** | | |
| $T1_{H_2O,\,GM}$ | 1.5 s | [24,36–40] |
| $T1_{H_2O,\,WM}$ | 0.97 s | [24,36–40] |
| $T1_{H_2O,\,CSF}$ | 4.47 s | [38] |
| $T2_{H_2O,\,GM}$ | 0.088 s | [24,39,41,42] |
| $T2_{H_2O,\,WM}$ | 0.073 s | [24,39,41,42] |
| $T2_{H_2O,\,CSF}$ | 2.030 s | [43] |
| **Metabolite** | | Metabolite relaxation parameters adopted in FSL-MRS are derived from an average of NAA, Cr and Cho values. |
| $T1_{Met}$ | 1.29 s | [34,37,44,45] |
| $T2_{Met}$ | 0.194 s | [29,34,44,45(p201),46,47] |
| MM | 1 | MM and $\kappa$ factors not considered in FSL-MRS |
| $\kappa$ | 1 | MM and $\kappa$ factors not considered in FSL-MRS |

*Supplementary Table 4 Relaxation parameters used for water scaling in FSL-MRS (all other parameters are per Supplementary Table 3)*

#### D.2 Gannet

Gannet models the water peak as a Lorentzian multiplied by a Gaussian at the same centre frequency, incorporating a linear baseline into the model. The water area is taken as the sum of that optimized function over the 3.8 - 5.6 ppm range (see https://github.com/richardedden/Gannet3.1/blob/master/GannetFit.m#L743)

Gannet can report a number of different water-scaled estimates, with the possibility for tissue-class correction per [48] and normalisation per [20], with integrated masking and segmentation functionality. The present study does not make use of these features, instead basing findings on metab.ConcIU field, which is calculated equivalently to Equation 2, with [$H_2O$] = 55000 mM and all other parameters per Supplementary Table 3. See also https://github.com/richardedden/Gannet3.1/blob/master/GannetFit.m#L1546

##### D.3 jMRUI: AMARES

Although jMRUI provides basic water referencing mechanisms internally, based on the amplitude of the reference FID, this functionality could not be batched effectively. Therefore, a separate AMARES run was used to model the water reference, using a superposition of one Gaussian and one Lorentzian component to mimic a pseudo-Voigt model. The centre frequency for the two components was constrained to be equal, with starting values for frequency and linewidth of 4.65 ppm and 7 Hz respectively; zero-order phase and begin time were set to zero.

Intrinsic amplitudes are reported for both the water reference and the metabolite models (i.e., $S_M$ and $S_{H2O}$ terms are available directly), hence there is no need to reverse any applied scaling in this case.

##### D.4 jMRUI: QUEST

Water referencing is performed identically to the AMARES case (D.3), using the same values for water amplitude.

##### D.5 LCModel

LCModel measures area of the water peak by integration: it first locates the maximum value of the frequency-domain water reference data within the expected range (4.65 +/- 1 ppm), then integrates the phase-corrected water reference +/-2 ppm around that maximum, assuming a linear baseline between the border regions. Refer LCModel.f lines 546, 4940, 5063 (available via http://www.s-provencher.com/lcmodel.shtml).

LCModel applies a scaling factor to the data, such that the signal strength per proton resonance is consistent between the data and the basis set (see also the LCModel user manual, http://www.s-provencher.com/pub/LCModel/manual/manual.pdf section 10.2):

$$f_{scale} = {Basis_{norm}}/{Water_{norm}}$$

$$Basis_{norm} = {S_{\mathrm{M}}}/{[\#H_{\mathrm{Met}} \cdot Conc_{Met} \cdot ATTMET]}$$

$$Water_{norm} = {S_{\mathrm{H_2O}}}/{[\#H_{\mathrm{H_2O}} \cdot WCONC \cdot ATTH2O]}$$

In this case, metabolite parameters are for NAA, the metabolite specified for basis set scaling (see C.5); $Conc_{Met}$. and $ATTMET$ are appropriate to the basis set (both 1.0), $\#H_{\mathrm{Met}}$ = 3 for the 2.01 ppm NAA peak. Given a consistently-scaled basis set, this is equivalent to scaling of the form:

$$\frac{S_{\mathrm{M}}}{S_{\mathrm{H_2O}}} \cdot \#H_{\mathrm{H_2O}} \cdot WCONC \cdot ATTH2O$$

ATTH2O accounts for the attenuation of NMR-visible water signal due to additional relaxation effects, approximated as $\exp\left(\frac{-\mathrm{TE}}{\mathrm{T2_{H_2O}}}\right)$. WCONC specifies the NMR-visible water concentration ($[\mathrm{H_2O}] \cdot Vis_{\mathrm{H_2O}}$). Default values of ATTH2O = 0.43 (for TE = 68 ms MEGA-PRESS, see section C.5) and WCONC = 35880 (pure WM) are adopted.

#### D.6 Osprey

Osprey uses a simulated water basis function to model the water reference, on a fit range from 2-7.4 ppm.

While the standardised processing pipeline evaluates the difference spectrum as diff = ( edit-OFF - edit-ON ), Osprey assumes a different convention for this: (edit-OFF - edit-ON) / 2. Therefore, an additional factor-of-two correction is needed in this instance.

Raw water scaled estimates (without tissue correction) are obtained. These are calculated similarly to Equation 2, but without macromolecule and edit efficiency terms:

$$\frac{S_{\mathrm{M}}}{S_{\mathrm{H_2O}}} \cdot [\mathrm{H_2O}] \cdot Vis_{\mathrm{H_2O}} \cdot \frac{\#H_{\mathrm{H_2O}}}{\#H_{\mathrm{Met}}} \cdot \frac{1-\exp\left(\frac{-\mathrm{TR}}{\mathrm{T1_{H_2O}}}\right)}{1-\exp\left(\frac{-\mathrm{TR}}{\mathrm{T1_{Met}}}\right)} \cdot \frac{\exp\left(\frac{-\mathrm{TE}}{\mathrm{T2_{H_2O}}}\right)}{\exp\left(\frac{-\mathrm{TE}}{\mathrm{T2_{Met}}}\right)}$$

In this scaling, Osprey assumes $[\mathrm{H_2O}]$ = 55500 mM, and slightly different parameters for Glx and tCr, including updated $\mathrm{T2_{Met}}$ estimates from [47]; relaxation estimates in this mode are averaged across tissue class and constituent metabolites (Glu + Gln for Glx, Cr + PCr for tCr). All other parameters are in accordance with Supplementary Table 3.

| **Parameter** | **Value** | **Source and Remarks** |
|---|---|---|
| **Water** | | |

| | | |
|---|---|---|
| $[H_2O]$ | 55500 mM | |
| **Glx** | | |
| $T1_{Met}$ | 1.265 s | [32,34] |
| $T2_{Met}$ | 0.133 s | [47] |
| **Cr** | | |
| $T1_{Met}$ | 1.350 s | [34] |
| $T2_{Met}$ | 0.156 s | [47] |

*Supplementary Table 5: Relaxation parameters used for water scaling in Osprey (all other parameters per Supplementary Table 3)*

See also: https://github.com/schorschinho/osprey/blob/develop/quantify/OspreyQuantify.m#L492

#### D.7 Tarquin

Tarquin uses the infinity norm (peak absolute amplitude) of the time-domain water reference data for normalisation to water. The reported value is given as:

$$\frac{S_M}{S_{\text{H2O}}} \cdot \#H_{\text{H2O}} \cdot \text{WCONC} \cdot \text{WATT}$$

Where WCONC represents the NMR-visible water concentration (equivalent to $[\text{H}_2\text{O}] \cdot Vis_{\text{H}_2\text{O}}$), and WATT accounts for the reduction of measured water signal relative to metabolite signal due to differences in T2 relaxation (equivalent to $\frac{\exp\left(\frac{-\text{TE}}{\text{T2}_{\text{H}_2\text{O}}}\right)}{\exp\left(\frac{-\text{TE}}{\text{T2}_{\text{Met}}}\right)}$); in this study, the WCONC and WATT parameters are specified as 35880 mM (pure white-matter value) and 0.76 respectively, noting that the latter is unlikely to be optimal for TE = 68 ms. Internally, $S_M$ is pre-scaled by a factor of 0.5 to account for internal basis-set scaling conventions. Refer: https://github.com/martin3141/tarquin/blob/master/src/common/Workspace.hpp#L469

### E Creatine Reference: Edit-OFF sub-spectrum modelling

For modelling of the edit-OFF sub-spectra by basis set algorithms (FSL-MRS, QUEST, LCModel, Osprey and Tarquin), a standard simulated basis set specific to each hardware vendor was adopted. These were derived for edit-OFF sub-spectra using a similar method to that used for the difference spectra (as detailed in section 2.3.1), incorporating a default set of basis components as defined within Osprey: ascorbate (Asc), aspartate (Asp), creatine (Cr), a negative correction term for the Creatine $CH_2$ singlet around 3.94 ppm (CrCH2), GABA, glycerophosphocholine (GPC), GSH Gln Glu, $H_2O$, myo-inositol (Ins), lactate (Lac), NAA, NAAG, phosphocholine (PCh), phosphocreatine (PCr), phosphoethanolamine (PE), scyllo-inositol (Scyllo), taurine (Tau), tyrosine (Tyros), and a series of macromolecule and lipid components: $MM_{0.9}$ $MM_{1.2}$ $MM_{1.4}$ $MM_{1.7}$ $MM_{2.0}$ $Lip_{0.9}$ $Lip_{1.3}$ $Lip_{2.0}$.

#### E.1 FSL-MRS

Edit-OFF sub-spectra were quantified with the standard edit-OFF basis set, using Cr+PCr as an internal reference for basis set scaling. For expedience, the default Newton optimisation algorithm was used; otherwise, all parameters were the same as for the difference case.

```
--basis             ge_off_8192_4000.basis
--data              S12_GABA_68.off.nii.gz
--h2o               S12_GABA_68.ref.nii.gz
--output            S12_GABA_68.off
--overwrite
--TE                68
--TR                2.0
--tissue_frac       .5 .5 0
--verbose
--report
--combine           NAA NAAG
--combine           Glu Gln GSH
--combine           Cr PCr
--combine           GPC PCh

Defaults:
--internal_ref      Cr PCr
--algo Newton
--ppmlim:           (0.2, 4.2)
--baseline_order:   2
--metab_groups:     0
--h2o_scale:        1.0
```

#### E.2 Gannet

In addition to the GABA+Glx model, Gannet independently fits a Lorentzian model to NAA around 2.01 ± 0.04 ppm, and a dual-Lorentzian model for Choline and Creatine, the latter centred at 3.02 ppm with a fixed 0.18 ppm separation, all from the edit-OFF sub-spectrum. A minor local modification was made to additionally yield water-referenced estimates from the existing Cr and NAA fits (usually only reported for those metabolites contained in the difference spectrum).

#### E.3 jMRUI: AMARES

For the edit-OFF sub-spectra, a simple model including only three major peaks and residual water was applied: NAA (2.0 ± 0.05 ppm), Cr (3.0 ± 0.05 ppm), Cho (3.2 ± 0.05 ppm), residual water (4.62 ± 0.08 ppm). Soft constraints on linewidth (2-15 Hz for metabolite, 4-25 Hz for water) and relative phase (±10 degrees for metabolites; unconstrained for water). As with the difference case, zeroth- and first-order phase were set to zero.

#### E.4 jMRUI: QUEST

The standard per-manufacturer edit-OFF basis sets described above were imported in jMRUI text format, manually aligned for NAA @ 2.0 ppm, before assembly into a jMRUI-format metabolite list. Other model parameters were the same as for the difference spectra.

#### E.5 LCModel

Edit-OFF sub-spectra were modelled with the standard per-manufacturer edit-OFF basis sets. Basic configuration was similar to that used for the difference spectra, with two key exceptions. `PPMGAP` was not defined – which implies fitting over the full defined fit range (0.2 - 4.2 ppm), the default mode of operation. `SPTYPE` was also not defined, again implying a default fit: a regular spline baseline was modelled (with default knot spacing 0.15 ppm), Creatine was used for internal basis set scaling, and LCModel's default set of soft constraints were adopted.

#### E.6 Osprey

Edit-OFF sub-spectra were modelled using the standard per-manufacturer edit-OFF basis sets, with otherwise similar parameters to those used for the difference spectra. This happened in the same invocation of OspreyFit.

#### E.7 Tarquin

Edit-OFF sub-spectra were quantified with the standard per-manufacturer edit-OFF basis sets, with HSVD water removal disabled (`--water_width 0`) but all other parameters equivalent to those used for the difference spectra. Tarquin (internal) used `--int_basis 1h_brain`.

```
tarquin

--format      lcm
--input       S12_GABA_68.off.RAW
--input_w     S12_GABA_68.off.REF

--echo        0.068
--fs          5000
--ft          127714400
--auto_phase  false
--auto_ref    false
--water_eddy  false
--water_width 0

--w_att       0.76
--w_conc      35880

--output_txt  S12_GABA_68.off.txt
--output_csv  S12_GABA_68.off.csv
--output_pdf  S12_GABA_68.off.pdf
--output_fit  S12_GABA_68.off.fit
--ext_pdf     true
```

For “Tarquin” (with standard basis set):

```
--basis_lcm   ge_off_4096_5000.basis
```

Or for “Tarquin (internal)” (internal basis set):

```
--int_basis   1h_brain
```

### F Quality Control

a) R1: Preprocessing

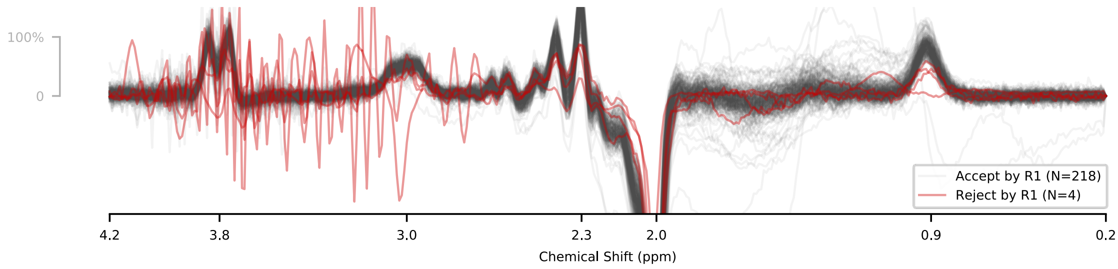


b) R2: SNR, Linewidth ($NAA_{diff}$)

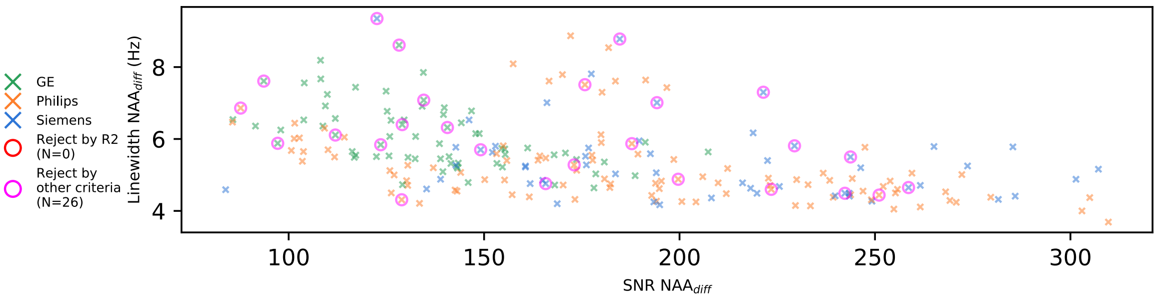


c) R3: Uncertainty (%SD, CRLB or %FitError)

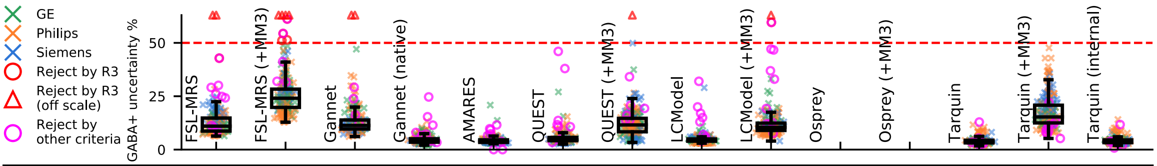


| Uncertainty Metric | CRLB | CRLB | FitError | FitError | SD | SD | SD | CRLB | CRLB | N/A | N/A | SD | SD | SD |
|---|---|---|---|---|---|---|---|---|---|---|---|---|---|---|
| GABA+ uncertainty % | 11±3.8 | 24±6.4 | 11±3 | 4.1±1.1 | 3.9±0.86 | 4.8±1.1 | 12±4.8 | 4±1.5 | 11±2.4 | - | - | 3.8±0.92 | 15±5.3 | 3.9±0.93 |
| Glx uncertainty % | 12±2.8 | 12±3.2 | 6.2±1.6 | 2.9±0.72 | 7.6±2.2 | 4.8±1.3 | 4.5±1.2 | 3±1.5 | 2±0 | - | - | 1.4±0.4 | 1.4±0.39 | 3.3±1.2 |
| tCr uncertainty % | 8.7±2.1 | 8.7±2.1 | 2.1±0.3 | 1.4±0.19 | 2.2±0.66 | 6.4±2.9 | 6.4±2.9 | 1±0 | 1±0 | - | - | 0.8±0.33 | 0.8±0.33 | 1±0.47 |
| Reject by R3 | 3 | 8 | 2 | 0 | 0 | 0 | 1 | 0 | 2 | 0 | 0 | 0 | 0 | 0 |

d) R4: Median Absolute Deviation (MAD) from global median

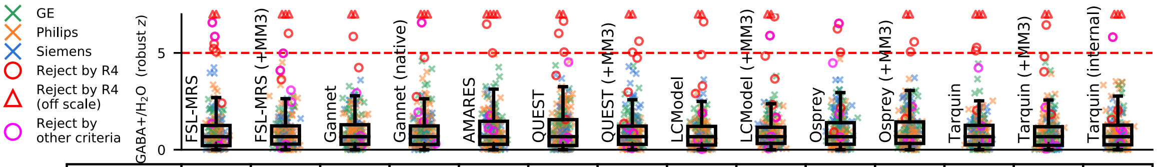


| | | | | | | | | | | | | | | |
|---|---|---|---|---|---|---|---|---|---|---|---|---|---|---|
| Reject by R4 | 9 | 7 | 4 | 6 | 6 | 7 | 8 | 7 | 8 | 5 | 5 | 6 | 6 | 6 |
| Invalid/Failed | 0 | 0 | 1 | 1 | 0 | 0 | 0 | 0 | 0 | 0 | 0 | 2 | 2 | 1 |
| TOTAL | 11 | 11 | 7 | 9 | 9 | 10 | 11 | 9 | 10 | 8 | 8 | 9 | 10 | 9 |
| Total GE | 2 (2%) | 3 (4%) | 2 (2%) | 4 (5%) | 2 (2%) | 3 (4%) | 4 (5%) | 2 (2%) | 2 (2%) | 2 (2%) | 4 (5%) | 3 (4%) | 2 (2%) | 2 (2%) |
| Total Philips | 2 (1%) | 1 (0%) | 3 (2%) | 2 (1%) | 1 (0%) | 3 (2%) | 3 (2%) | 3 (2%) | 3 (2%) | 2 (1%) | 1 (0%) | 2 (1%) | 4 (3%) | 2 (1%) |
| Total Siemens | 7 (12%) | 7 (12%) | 2 (3%) | 3 (5%) | 6 (11%) | 4 (7%) | 4 (7%) | 4 (7%) | 5 (9%) | 4 (7%) | 3 (5%) | 4 (7%) | 4 (7%) | 5 (9%) |

*Supplementary Figure 6: Quality control; rejected fits for each criterion, according to algorithm. A single dataset may be flagged by multiple rejection criteria.*

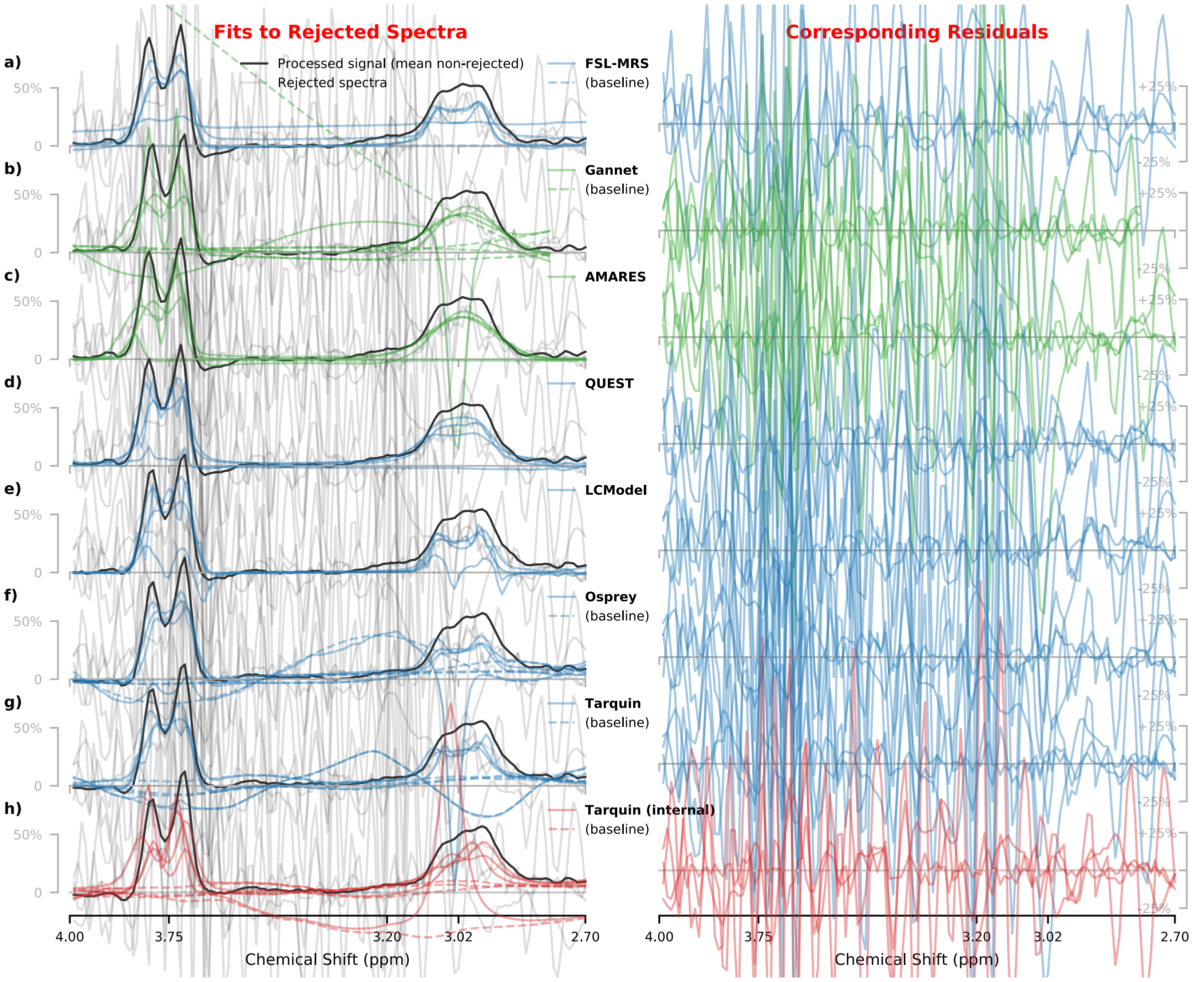


*Supplementary Figure 7: Most algorithms dutifully applied their model even when supplied with very poor input data, often returning visually pleasing fits which were acceptable according to other criteria.*

### G Fitting Outcomes per algorithm

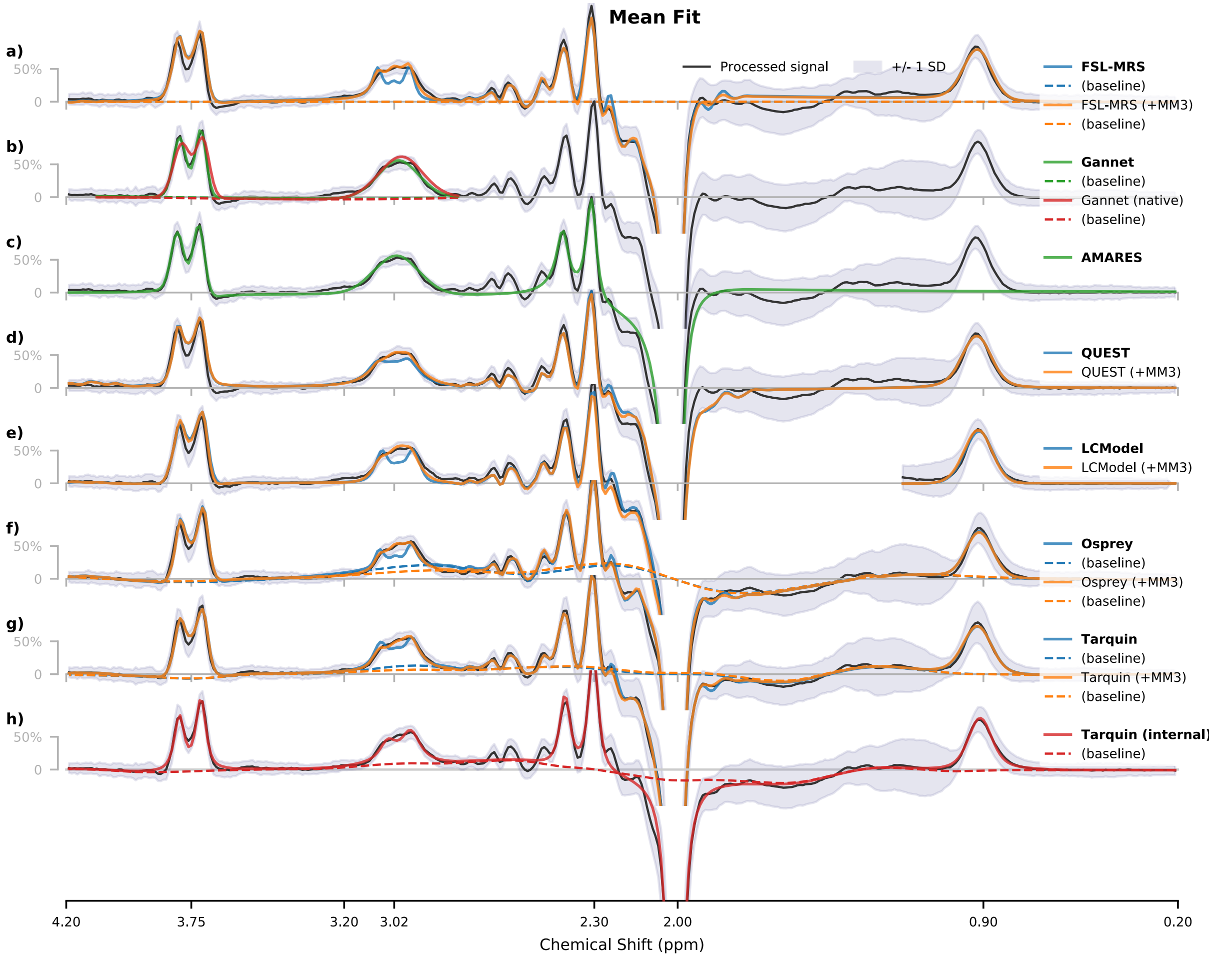


*Supplementary Figure 8: Average metabolite and baseline (where applicable) models with corresponding residuals for the GABA+ edited spectra, for each algorithm. Vertical scaling is normalised. This represents the same data as Figure 2, on the full fit range for each algorithm*

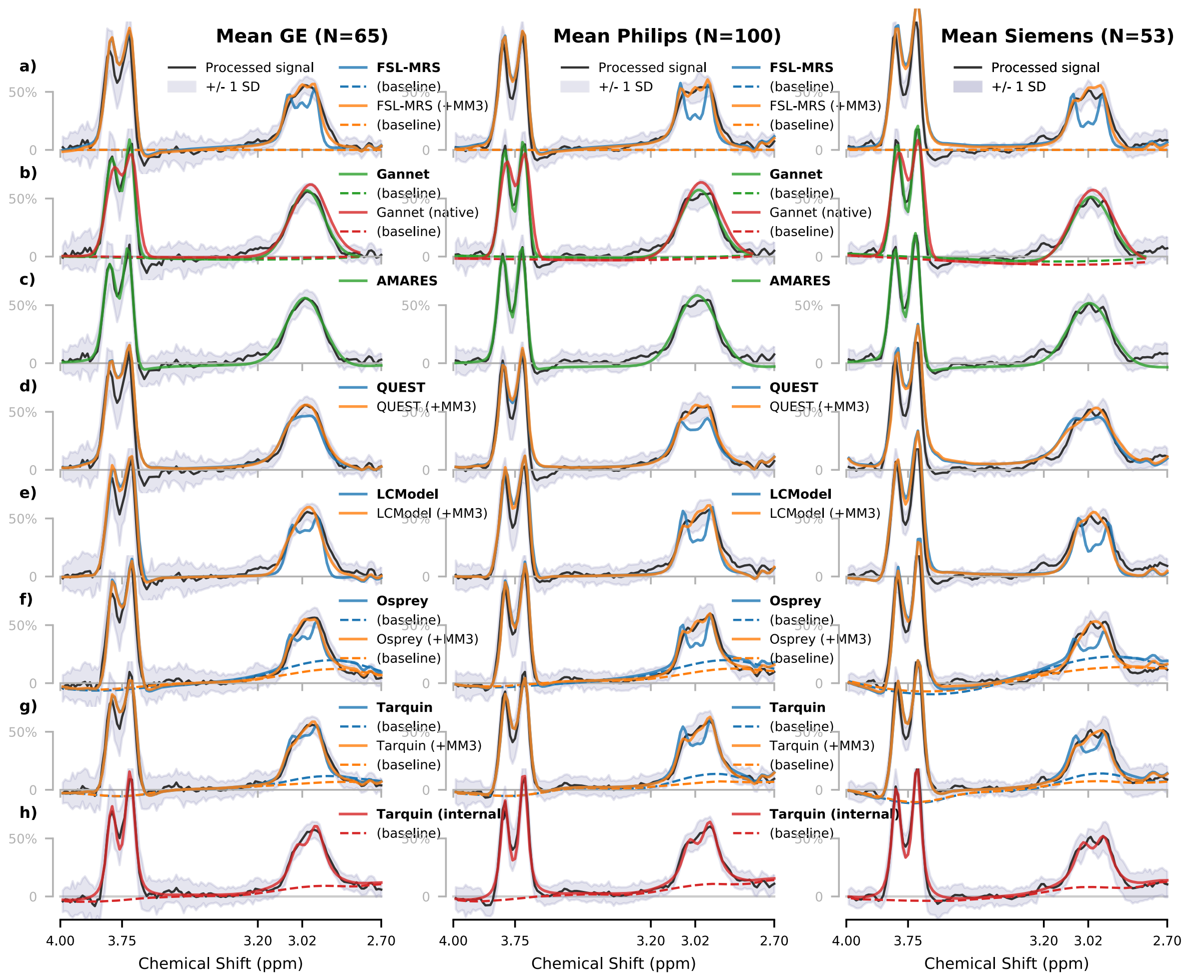


*Supplementary Figure 9: Average metabolite and baseline (where applicable) models for the GABA+ edited spectra, for each algorithm and each vendor. Vertical scaling is normalised. This represents the same data as Figure 2, split according to vendor.*

### H Agreement between algorithms

.

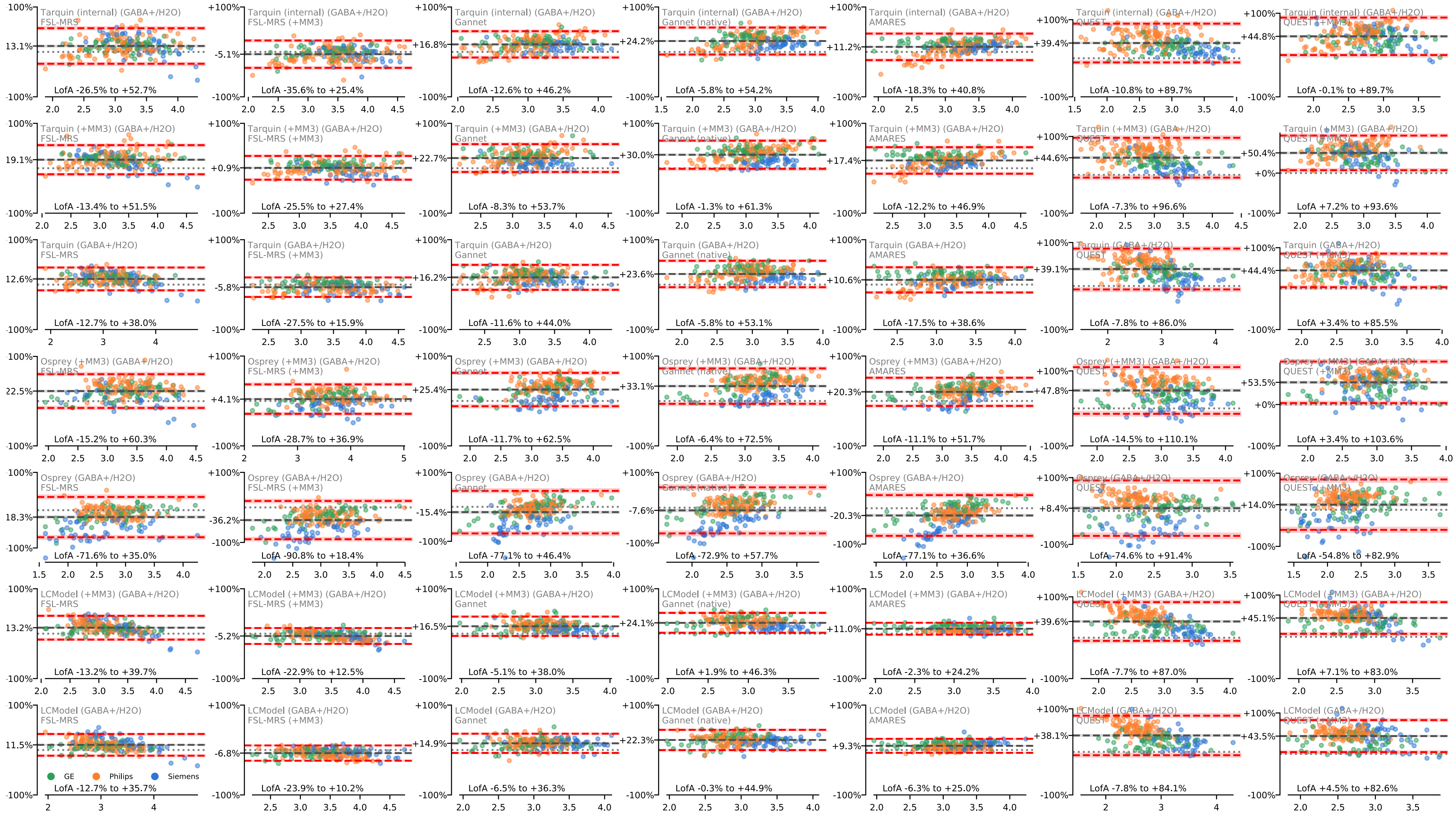


*Supplementary Figure 10: Bland-Altman plots, comparing pairs of estimates for GABA+/ $H_2O$ and GABA+/tCr (continued overleaf…)*

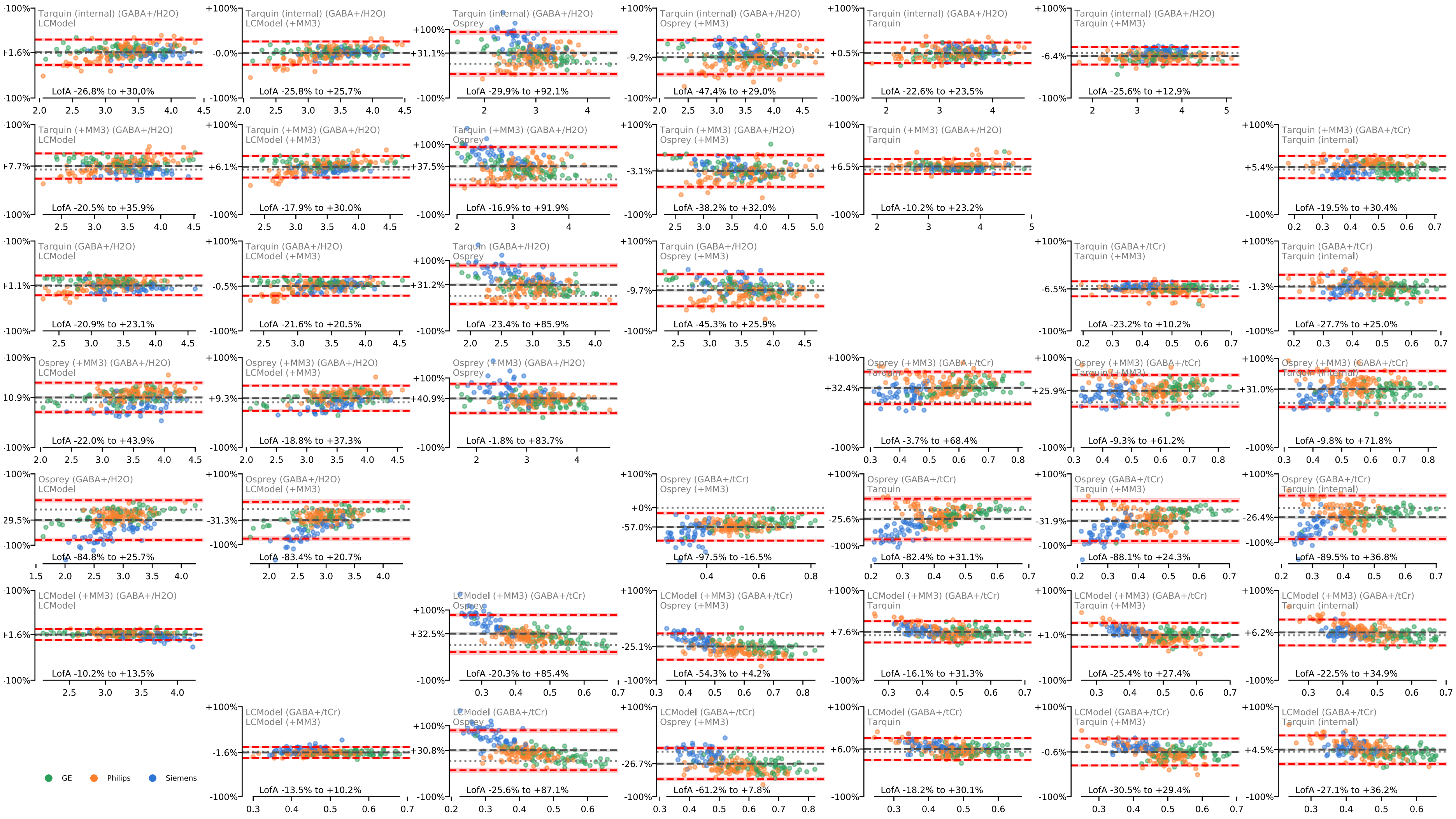


*Supplementary Figure 10 (…continued…)*

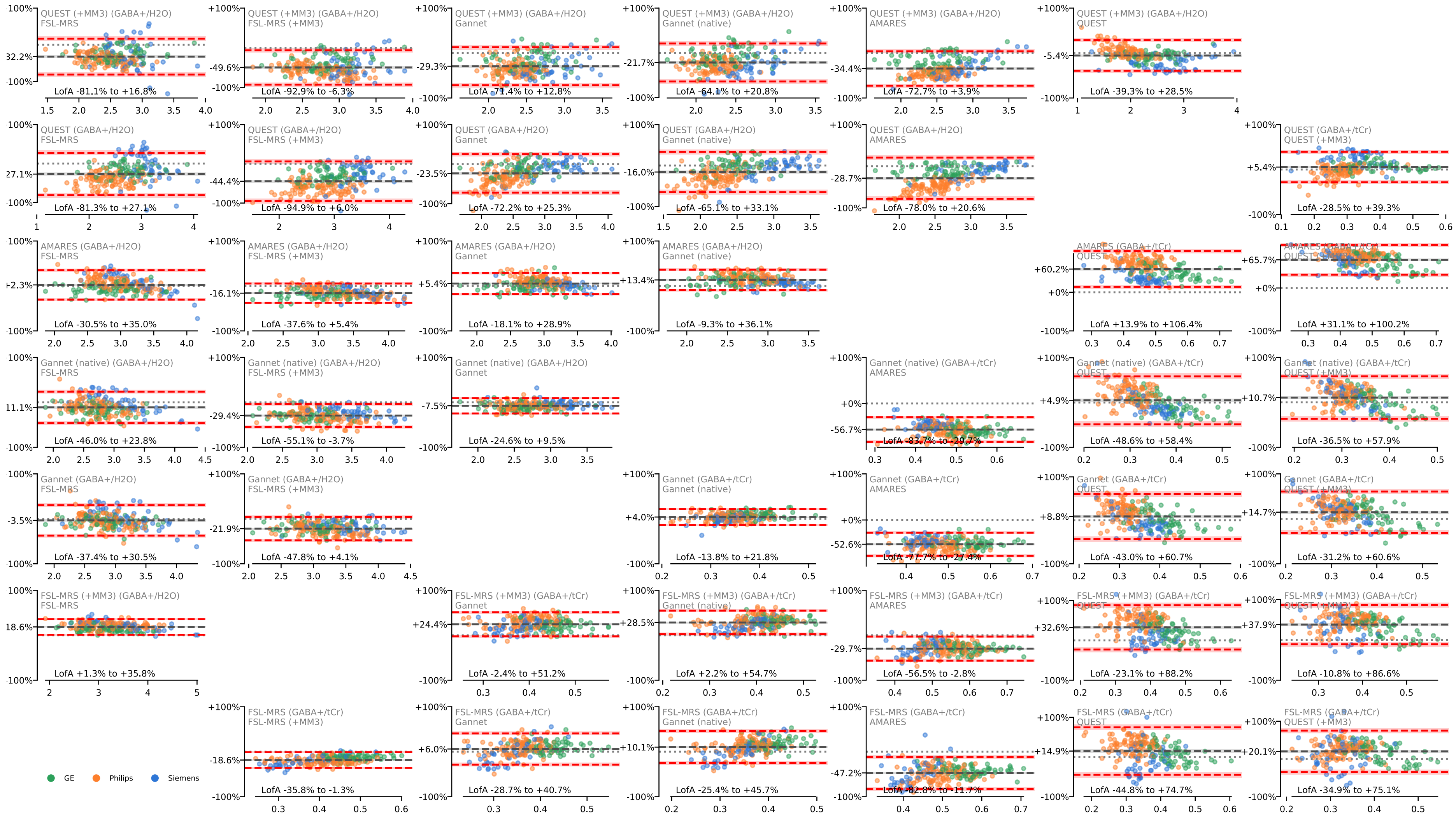


*Supplementary Figure 10 (…continued…)*

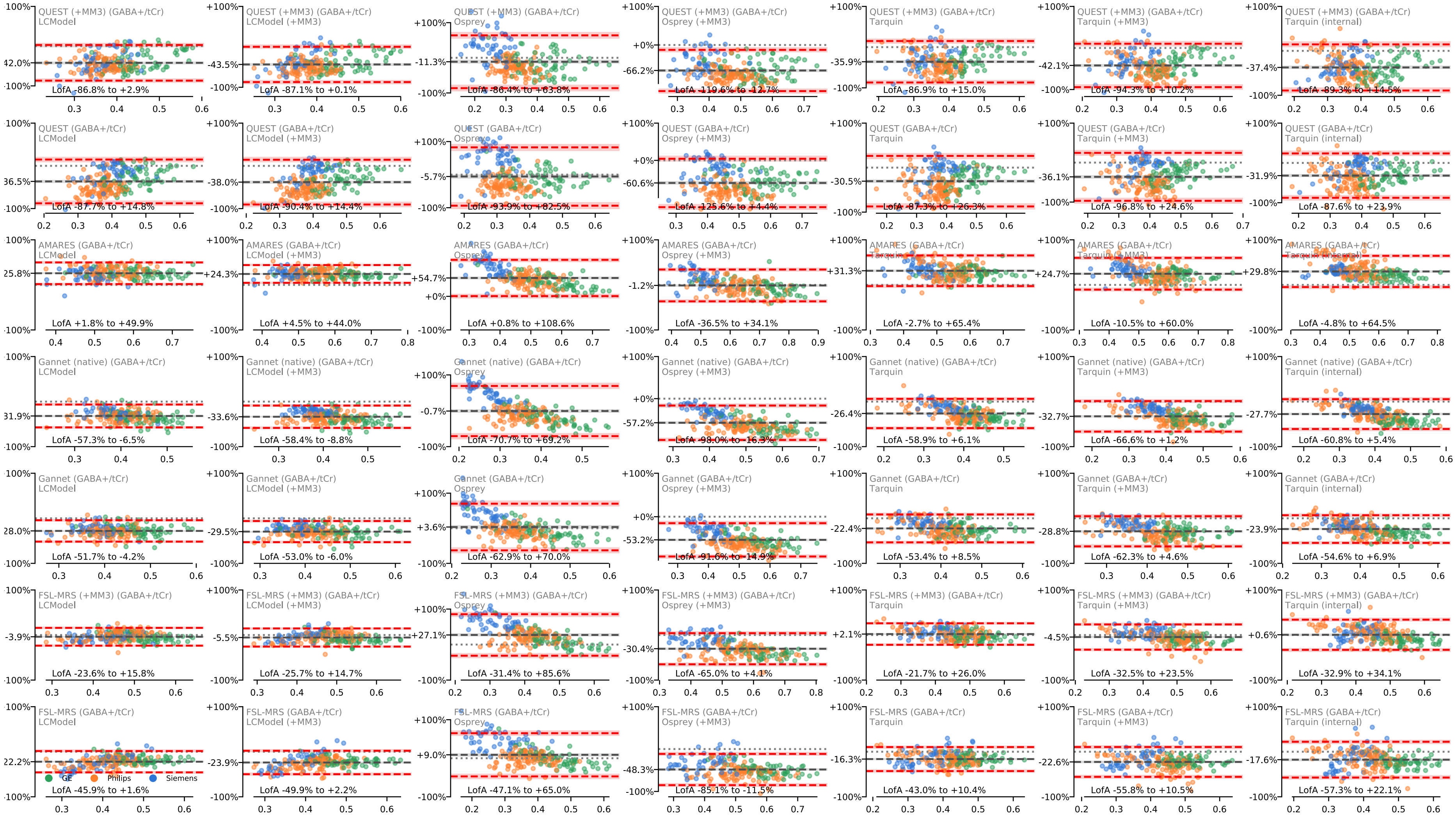


*Supplementary Figure 10 (…continued)*

*Supplementary Figure 11: Distribution of metabolite estimates obtained from each algorithm, grouped by vendor*

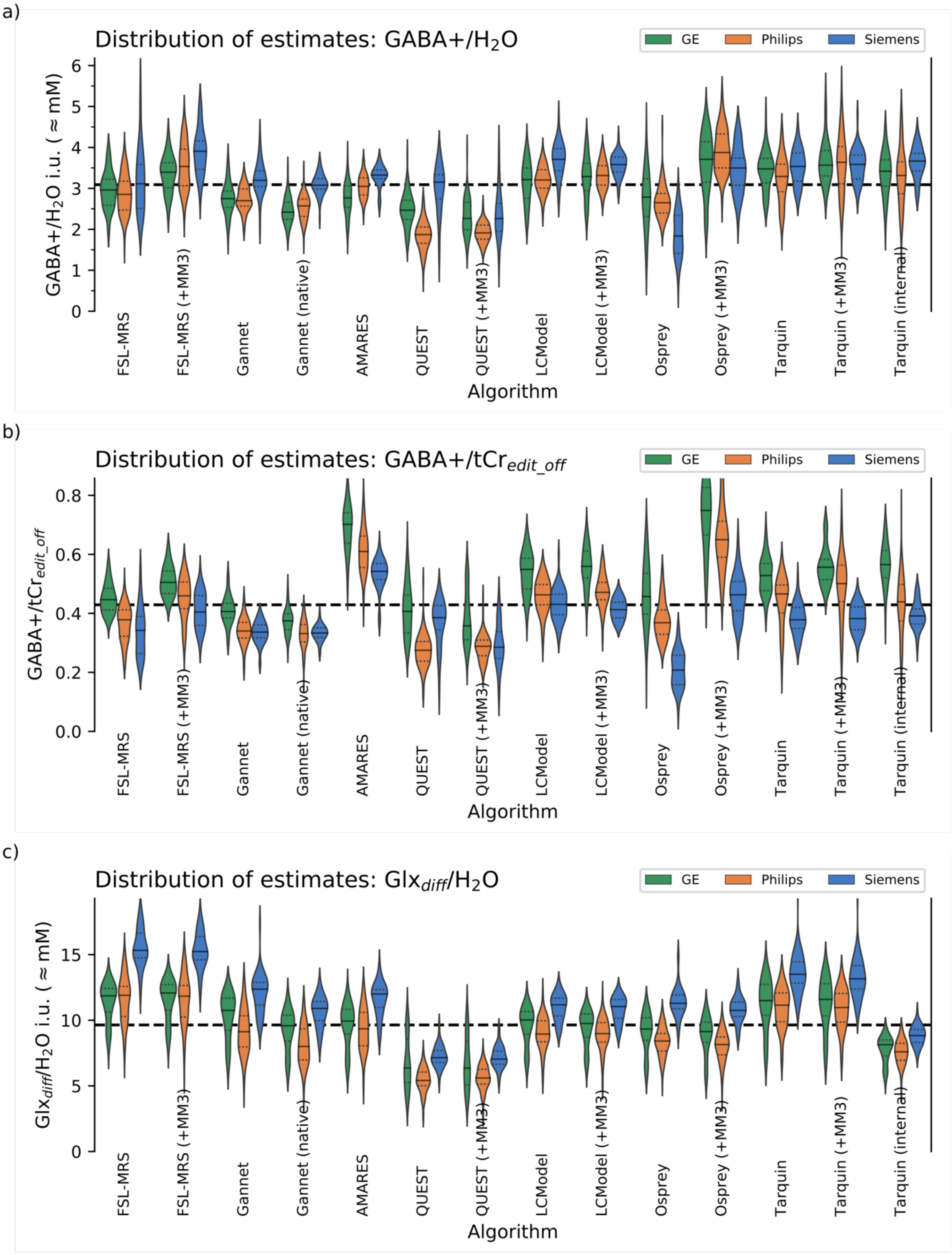

d)

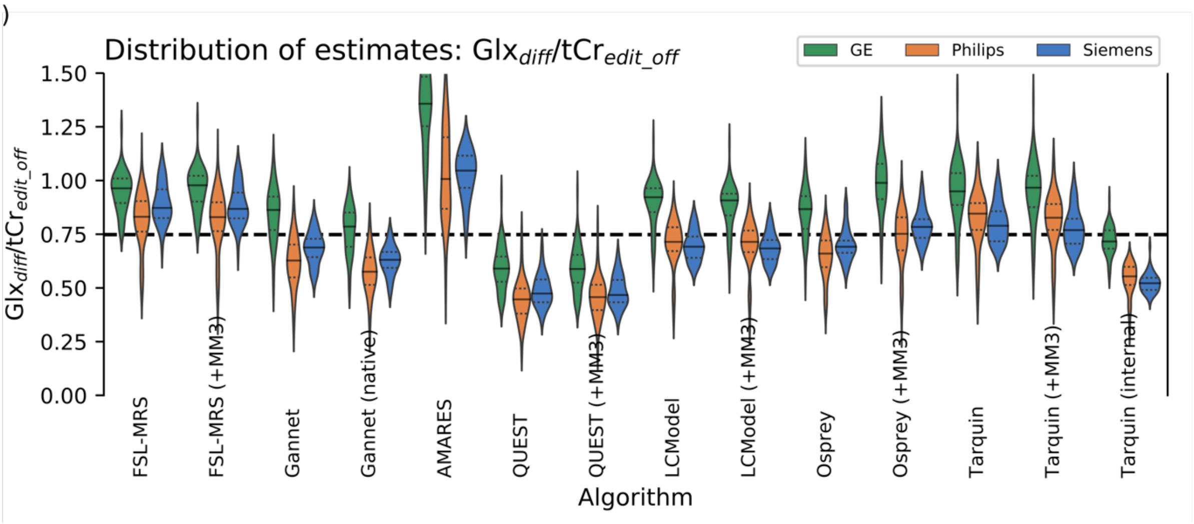
Distribution of estimates: $Glx_{diff}/tCr_{edit\_off}$
GE
Philips
Siemens
$Glx_{diff}/tCr_{edit\_off}$
1.50
1.25
1.00
0.75
0.50
0.25
0.00
FSL-MRS
FSL-MRS (+MM3)
Gannet
Gannet (native)
AMARES
QUEST
QUEST (+MM3)
LCModel
LCModel (+MM3)
Osprey
Osprey (+MM3)
Tarquin
Tarquin (+MM3)
Tarquin (internal)
Algorithm

e)

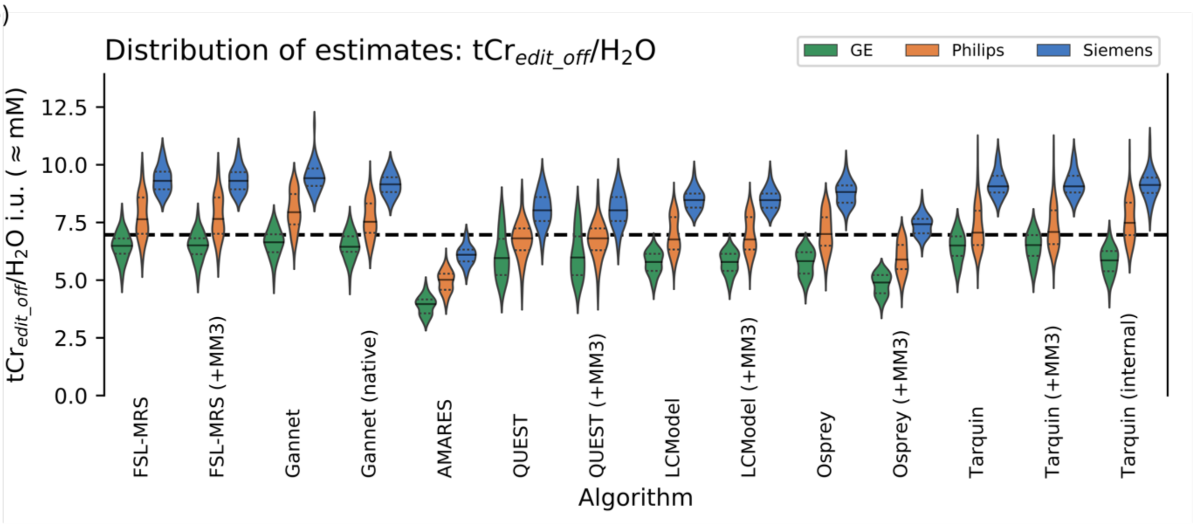
Distribution of estimates: $tCr_{edit\_off}/H_2O$
GE
Philips
Siemens
$tCr_{edit\_off}/H_2O$ i.u. ( ≈ mM)
12.5
10.0
7.5
5.0
2.5
0.0
FSL-MRS
FSL-MRS (+MM3)
Gannet
Gannet (native)
AMARES
QUEST
QUEST (+MM3)
LCModel
LCModel (+MM3)
Osprey
Osprey (+MM3)
Tarquin
Tarquin (+MM3)
Tarquin (internal)
Algorithm

| | FSL-MRS | FSL-MRS (+MM3) | Gannet | Gannet (native) | AMARES | QUEST | QUEST (+MM3) | LCModel | LCModel (+MM3) | Osprey | Osprey (+MM3) | Tarquin | Tarquin (+MM3) | Tarquin (internal) | Median |
|---|---|---|---|---|---|---|---|---|---|---|---|---|---|---|---|
| GABA+/H2O mean ±SD | 2.950 ±0.567 | 3.541 ±0.559 | 2.837 ±0.409 | 2.644 ±0.388 | 3.059 ±0.375 | 2.230 ±0.633 | 2.072 ±0.456 | 3.308 ±0.462 | 3.379 ±0.396 | 2.538 ±0.639 | 3.707 ±0.620 | 3.441 ±0.509 | 3.591 ±0.588 | 3.448 ±0.593 | 3.190 ±0.401 |
| GABA+/H2O diff GE | +0.4% | -4.1% | -3.1% | * -8.5% | *** -9.5% | +10.6% | +9.3% | -2.9% | -2.7% | +9.8% | +0.1% | +1.0% | -0.7% | -0.9% | -1.4% |
| GABA+/H2O diff Philips | -3.3% | -0.2% | -4.8% | -2.7% | -0.3% | *** -16.1% | *** -7.7% | -3.1% | -1.9% | +4.4% | +4.6% | -4.4% | +1.3% | -3.8% | -2.9% |
| GABA+/H2O diff Siemens | +5.6% | * +10.4% | *** +12.7% | *** +16.5% | *** +8.7% | *** +41.3% | +9.2% | *** +12.1% | ** +6.0% | *** -27.6% | -5.7% | +2.7% | -0.1% | +6.3% | ** +5.8% |
| GABA+/tCr mean ±SD | 0.393 ±0.076 | 0.466 ±0.072 | 0.358 ±0.055 | 0.341 ±0.046 | 0.611 ±0.092 | 0.321 ±0.094 | 0.301 ±0.081 | 0.477 ±0.074 | 0.471 ±0.077 | 0.366 ±0.123 | 0.639 ±0.146 | 0.466 ±0.090 | 0.497 ±0.104 | 0.458 ±0.106 | 0.441 ±0.076 |
| GABA+/tCr diff GE | *** +13.6% | *** +8.4% | *** +13.5% | *** +9.8% | *** +14.9% | *** +26.5% | *** +18.9% | *** +15.1% | *** +18.9% | *** +24.9% | *** +17.1% | *** +13.5% | *** +11.9% | *** +23.4% | *** +17.3% |
| GABA+/tCr diff Philips | -3.7% | -1.5% | -5.0% | -2.9% | -0.2% | *** -14.5% | ** -4.1% | -3.0% | +0.1% | +0.6% | +1.6% | +0.0% | +0.7% | -4.2% | -1.1% |
| GABA+/tCr diff Siemens | -12.8% | *** -13.2% | * -6.0% | -2.3% | *** -11.2% | +19.8% | -5.2% | *** -9.7% | *** -12.3% | *** -43.2% | *** -27.6% | *** -18.8% | *** -23.2% | *** -14.6% | *** -14.3% |
| Glx/H2O mean ±SD | 12.299 ±2.289 | 12.410 ±2.251 | 10.352 ±2.009 | 9.256 ±1.749 | 9.980 ±1.781 | 6.167 ±1.526 | 6.279 ±1.473 | 9.868 ±1.464 | 9.663 ±1.414 | 9.244 ±1.715 | 8.881 ±1.633 | 11.783 ±1.978 | 11.683 ±1.914 | 8.124 ±1.039 | 9.847 ±1.610 |
| Glx/H2O diff GE | ** -3.6% | -2.7% | +3.8% | +3.5% | -0.4% | +3.2% | +1.2% | +1.6% | +1.0% | +1.0% | +2.9% | -2.4% | -0.8% | +0.1% | +1.2% |
| Glx/H2O diff Philips | -3.2% | -4.6% | *** -11.8% | *** -13.5% | -6.3% | *** -12.3% | *** -10.8% | * -9.3% | * -7.1% | *** -8.9% | *** -8.2% | * -5.4% | -6.1% | -6.5% | ** -10.1% |
| Glx/H2O diff Siemens | *** +24.6% | *** +22.8% | *** +19.5% | *** +17.8% | *** +20.2% | *** +15.9% | *** +12.1% | *** +13.3% | *** +14.2% | *** +22.1% | *** +21.1% | *** +14.6% | *** +12.6% | *** +8.7% | *** +15.7% |
| Glx/tCr mean ±SD | 0.883 ±0.123 | 0.880 ±0.130 | 0.698 ±0.145 | 0.640 ±0.127 | 1.102 ±0.236 | 0.485 ±0.113 | 0.491 ±0.110 | 0.746 ±0.134 | 0.737 ±0.129 | 0.699 ±0.133 | 0.805 ±0.161 | 0.859 ±0.139 | 0.850 ±0.141 | 0.575 ±0.107 | 0.749 ±0.134 |
| Glx/tCr diff GE | *** +9.2% | *** +11.2% | *** +23.7% | *** +22.8% | *** +23.2% | *** +21.8% | *** +19.8% | *** +23.6% | *** +23.2% | *** +24.0% | *** +22.9% | *** +10.6% | *** +13.7% | *** +24.7% | *** +21.9% |
| Glx/tCr diff Philips | * -5.8% | * -5.7% | *** -10.0% | *** -10.0% | -8.6% | ** -7.8% | * -6.9% | * -4.3% | -3.1% | *** -5.6% | *** -6.6% | -1.5% | -2.8% | *** -3.6% | * -5.0% |
| Glx/tCr diff Siemens | -1.2% | -1.3% | -1.3% | -1.4% | ** -5.1% | -2.2% | -4.8% | *** -7.3% | *** -7.2% | -1.1% | -2.6% | -8.1% | ** -9.5% | *** -9.1% | * -6.7% |
| tCr/H2O mean ±SD | 7.368 ±1.287 | 7.370 ±1.276 | 7.794 ±1.267 | 7.496 ±1.175 | 4.814 ±0.897 | 6.838 ±1.114 | 6.845 ±1.114 | 6.652 ±1.189 | 6.634 ±1.185 | 6.719 ±1.291 | 5.694 ±1.089 | 7.098 ±1.281 | 7.109 ±1.278 | 7.275 ±1.412 | 7.072 ±1.223 |
| tCr/H2O diff GE | *** -12.0% | *** -11.7% | *** -14.8% | *** -14.0% | *** -17.7% | *** -12.9% | *** -12.6% | *** -13.0% | *** -12.8% | *** -13.3% | *** -14.0% | *** -8.4% | *** -8.3% | *** -19.5% | *** -13.9% |
| tCr/H2O diff Philips | +3.7% | +3.8% | +1.9% | +0.4% | +4.2% | -0.4% | -0.5% | +1.7% | +2.0% | +4.2% | +3.5% | -0.6% | -0.3% | +2.9% | +1.1% |
| tCr/H2O diff Siemens | *** +26.3% | *** +26.2% | *** +20.8% | *** +22.1% | *** +26.7% | *** +17.4% | *** +17.3% | *** +27.3% | *** +27.6% | *** +31.3% | *** +30.4% | *** +27.7% | *** +27.5% | *** +25.4% | *** +24.4% |

*Supplementary Table 6: Mean concentration estimates (institutional units, ≈ mM) from each algorithm, reported across all subjects. Estimates grouped by vendor are expressed as a % difference relative to the mean across all subjects for the respective algorithm; significance indicated by *, **, *** for $p_{holm}$ < .05, .01 and .001 respectively.*

### I Additional Correlation Analyses

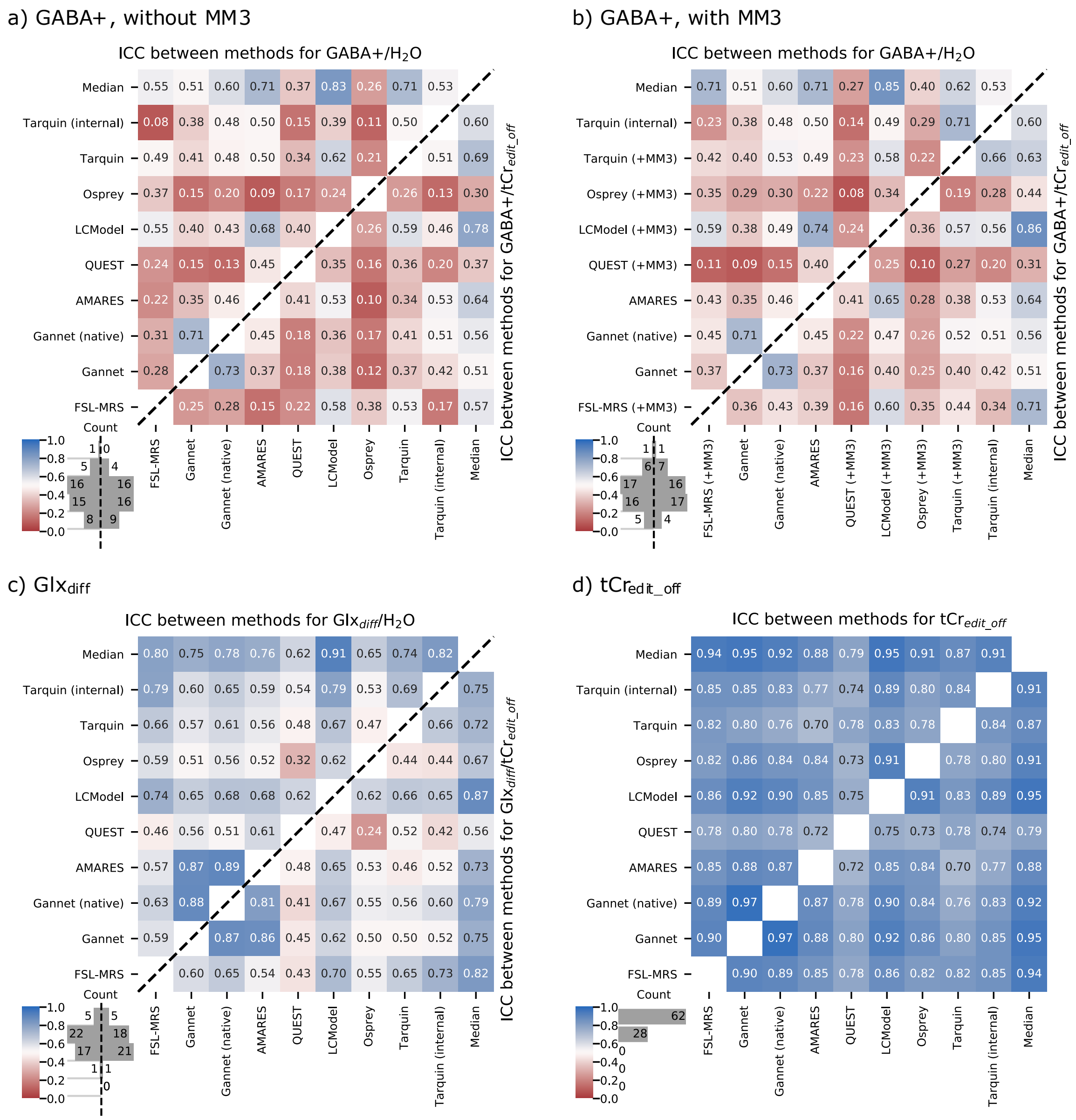


*Supplementary Figure 12 Intra-class correlation between algorithms, for additional metabolites and ratios*

#### I.1 $Glx_{diff}$, $Glx_{edit\_off}$

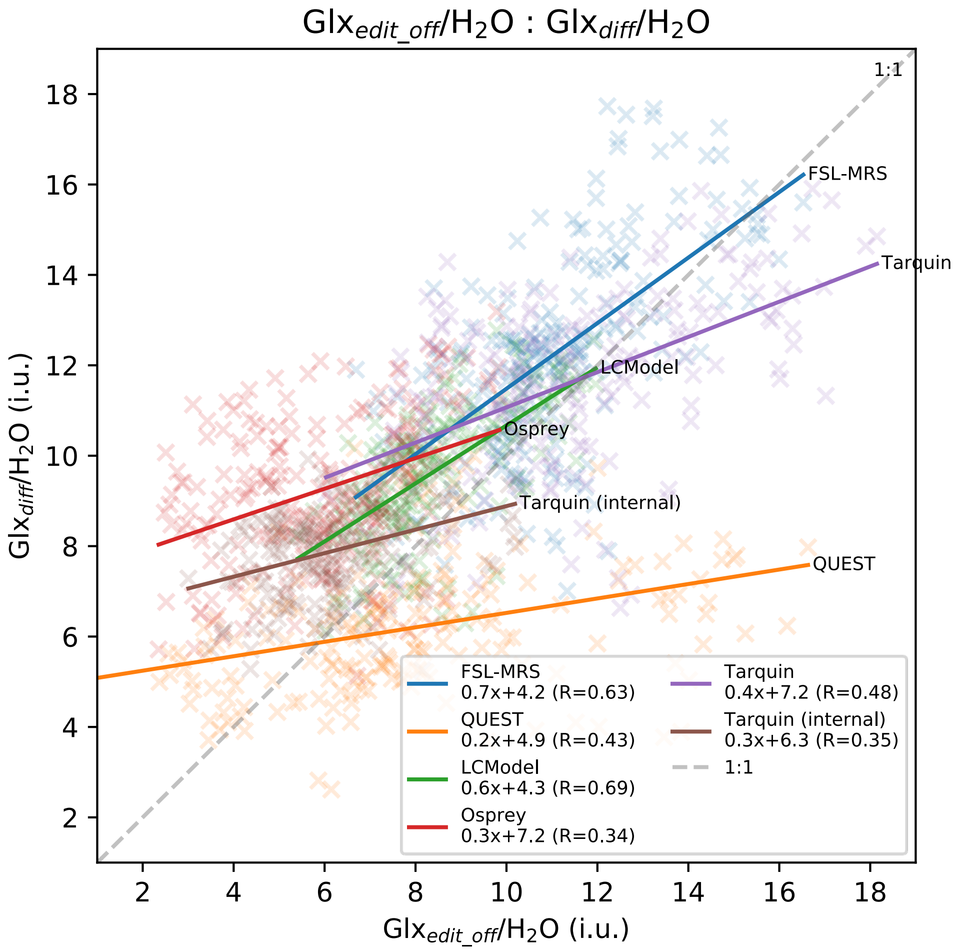


*Supplementary Figure 13: Estimates for $Glx_{edit\_off}$ vs $Glx_{diff}$ for each algorithm which modelled both.*

### K Figures Captions

Figure 1: Processing (b) and modelling (c) workflow, summarising key differences between the algorithms assessed.

Figure 2 Average metabolite and baseline (where applicable) models with corresponding residuals for the GABA+ edited spectra, for each algorithm. Vertical scaling is normalised; outcomes over the full fit range are presented in Supplementary Figure 8; outcomes split by vendor are presented in Supplementary Figure 9.

Figure 3: Distribution of GABA+/$H_2O$ estimates from each algorithm, grouped by manufacturer. Global median is shown in dashed black.

Figure 4 Relationship between GABA+ and grey matter, with different modelling strategies for GABA+. Robust (skipped) correlation coefficients are reported, with line-of-best-fit in dashed black.

Figure 5 Intraclass correlation coefficients between algorithms, scaled to water (upper left triangle) and $tCr_{edit\_off}$ (lower right triangle), with basis set algorithms excluding (a) and including (b) a component representing co-edited macromolecule contribution. "Median" data denotes correlation with the median estimate across all algorithms.

Supplementary Figure 1: Average metabolite and baseline (where applicable) models with corresponding residuals for each algorithm, baseline model and constraint model in the exploratory analysis. Corresponding fits over the full range are presented in Supplementary Figure 2.

Supplementary Figure 2: Average metabolite and baseline (where applicable) models for each algorithm, baseline model and constraint model in the exploratory analysis.

Supplementary Figure 3: Distribution of metabolite estimates for GABA+ (a), GABA (b), and MM3co (c) obtained from each modelling strategy, grouped by vendor

Supplementary Figure 4: Relationship between GABA+ and grey matter, with different baseline and soft constraint parameters for the MM3 component. Robust (skipped) correlation coefficients are reported, with line-of-best-fit in dashed black

Supplementary Figure 5: Relationship between GABA and grey matter, with different baseline and soft constraint parameters for the MM3 component. Robust (skipped) correlation coefficients are reported, with line-of-best-fit in dashed black

Supplementary Figure 6: Quality control; rejected fits for each criterion, according to algorithm. A single dataset may be flagged by multiple rejection criteria.

Supplementary Figure 7: Most algorithms dutifully applied their model even when supplied with very poor input data, often returning visually pleasing fits which were acceptable according to other criteria.

Supplementary Figure 8: Average metabolite and baseline (where applicable) models with corresponding residuals for the GABA+ edited spectra, for each algorithm. Vertical scaling is normalised. This represents the same data as Figure 2, on the full fit range for each algorithm

Supplementary Figure 9: Average metabolite and baseline (where applicable) models for the GABA+ edited spectra, for each algorithm and each vendor. Vertical scaling is normalised. This represents the same data as Figure 2, split according to vendor.

Supplementary Figure 10: Bland-Altman plots, comparing pairs of estimates for GABA+/ $H_2O$ and GABA+/tCr (continued overleaf…)

Supplementary Figure 11: Distribution of metabolite estimates obtained from each algorithm, grouped by vendor

Supplementary Figure 12 Intra-class correlation between algorithms, for additional metabolites and ratios

Supplementary Figure 13: Estimates for $Glx_{edit\_off}$ vs $Glx_{diff}$ for each algorithm which modelled both.

Supplementary Table 1: Basic demographics, hardware and software parameters for the constituent datasets

Supplementary Table 2: Mean concentration estimates (institutional units, ≈ mM), for each algorithm in the exploratory analysis. Estimates grouped by vendor are expressed as a % difference relative to the mean across all subjects for the respective algorithm; significance indicated by *, **, *** for $p_{holm}$ < .05, .01 and .001 respectively

Supplementary Table 3 Parameters for water-scaled metabolite estimates

Supplementary Table 4 Relaxation parameters used for water scaling in FSL-MRS (all other parameters are per Supplementary Table 3)

Supplementary Table 5: Relaxation parameters used for water scaling in Osprey (all other parameters per Supplementary Table 3)

Supplementary Table 6: Mean concentration estimates (institutional units, ≈ mM) from each algorithm, reported across all subjects. Estimates grouped by vendor are expressed as a % difference relative to the mean across all subjects for the respective algorithm; significance indicated by *, **, *** for $p_{holm}$ < .05, .01 and .001 respectively.
